# Towards an armed oncolytic virus approach to glioblastoma treatment

**DOI:** 10.1101/2025.03.07.641877

**Authors:** Arianna Calistri, Alberto Reale, Maria Vittoria Fornaini, Viola Donati, Ana Gabriela De Oliveira Do Rego, Mariateresa Panarelli, Alessandra Rossetto, Chiara Di Pietro, Marta Trevisan, Luca Persano, Elena Rampazzo, Daniela Marazziti, Fabio Mammano

## Abstract

Glioblastoma (GBM) is among the most aggressive and lethal human tumors. The current standard of care—surgical resection followed by chemotherapy—offers limited efficacy, as recurrence remains frequent and severe, underscoring the urgent need for novel therapeutic strategies. Photodynamic therapy (PDT) and oncolytic virotherapy have emerged as promising alternatives. PDT utilizes light-sensitive molecules to generate reactive oxygen species (ROS), selectively inducing tumor cell death, while oncolytic virotherapy employs viruses to lyse tumor cells and activate anti-tumor immune responses. Notably, Talimogene laherparepvec (T-VEC), an HSV-1-based oncolytic virus (oHSV1), is already approved for treating unresectable melanoma.

To explore a combinatorial approach for GBM, we engineered highly neuroattenuated oHSV1 variants with a genetic background similar to T-VEC, expressing KillerRed (KR)—a photosensitizing protein—alone or in combination with immunotherapeutic factors. Our results demonstrate potent cytolytic effects of these recombinant viruses in multiple murine and human GBM cell lines, as well as in primary tumor cells. In a syngeneic C57BL/6J mouse model, oHSV1 administration alone or carried by monocytes induced extensive tumor necrosis, accompanied by infiltration of CD3+ immune cells.

## Introduction

Glioblastoma (GBM) is the most common and aggressive brain tumor, with a poor prognosis despite current treatment options. Standard therapy, surgical resection followed by chemotherapy, often fails due to recurrence, highlighting the urgent need for novel therapeutic strategies [1, 2]. Among emerging approaches, oncolytic virotherapy and photodynamic therapy (PDT) are two promising strategies in cancer treatment, including GBM [3].

Oncolytic viruses (OVs) represent a versatile and promising tool for treating malignancies with poor prognoses [4]. These viruses exploit defects in cancer cells’ antiviral pathways, which also regulate proliferation. Beyond direct tumor lysis, OVs act as immunotherapies by eliciting an antitumor immune response and can be engineered to express therapeutic genes [5]. Most OVs are attenuated human viruses or wild-type viruses that naturally infect other species without causing disease in humans. Attenuation of human viruses minimizes their ability to evade antiviral defenses in healthy cells; for example, oncolytic herpes simplex virus type 1 (oHSV1) is often engineered with deletions in the γ34.5 gene to reduce neurovirulence [6]. The removal of both γ34.5 copies abolishes HSV-1 neurovirulence in animal models [7] and human patients [8], a critical safety consideration given that HSV-1 can cause encephalitis in immunocompetent adults.

T-VEC, an oHSV1 variant with deletions in γ34.5 and Us12 (a gene that reduces MHC-I antigen presentation), was the first OV approved by the FDA and EMA for clinical use, specifically for intratumoral injection in unresectable melanoma [9, 10]. T-VEC is also engineered to express granulocyte-macrophage colony-stimulating factor (GM-CSF) to enhance immune activation [11]. Several oHSV1 variants are currently in clinical trials for GBM treatment [12]. However, while early-phase trials showed promising results with intracranial oHSV1 injection, oncolytic virotherapy has yet to become a clinical reality for GBM. A key limitation is the delivery route: direct intracranial administration, typically via stereotactic injection or intrathecal catheterization, remains the only viable method [13–15]. These procedures, though feasible in research settings, are difficult to implement in large clinical trials and real-world practice. Moreover, except for intrathecal catheterization, these approaches do not easily permit repeated dosing of oHSV1.

To overcome these limitations, *ex vivo* loading of OVs into carrier cells has been proposed. These cells, once reinjected into the bloodstream, can exploit their natural tumor-homing properties to deliver OVs to the tumor site [16]. In the context of GBM, carrier cells must also be capable of crossing the blood-brain barrier (BBB). Monocytes, which constitute approximately 10% of circulating leukocytes, are promising candidates for this role. As precursors of tumor-associated macrophages (TAMs) in GBM [17], monocytes can infiltrate the central nervous system [18]. These cells can be isolated from peripheral blood, infected *ex vivo* with oHSV1, and then used as Trojan horses for systemic virus delivery. Despite their potential, monocytes have been scarcely explored as OV carriers, and never for oHSV1 [19, 20]. Notably, monocytes can be infected by wild-type HSV-1 but do not support high-level viral replication [21], a favorable characteristic since carrier cells must remain viable and motile until reaching the tumor. We recently demonstrated that oHSV1-infected human monocytes migrate toward breast cancer cells *in vitro* and head-and-neck squamous cancer cells in an *in ovo* chorioallantoic membrane (CAM) model [22, 23].

PDT, another promising therapy, employs a photosensitizing agent, light, and oxygen to selectively kill cancer cells [24]. The INDYGO clinical trial is currently evaluating intraoperative PDT following GBM resection using protoporphyrin IX (PpIX) induced by 5-aminolevulinic acid (5-ALA) [25]. Interstitial PDT (iPDT), involving stereotactic fiber optic placement for light delivery, has been experimentally applied to brain tumors for over 30 years [26]. However, limitations in light delivery and photosensitizer design have hindered the broader clinical adoption of PDT for GBM [25].

Genetically encoded photosensitizers, such as the KillerRed (KR) protein, offer an alternative approach. KR generates reactive oxygen species (ROS) upon irradiation with 540–590 nm light [27], making it suitable for combination therapies, including immunotherapy [28] and targeted photodynamic virotherapy [29]. A recent study has explored the potential of combining oHSV1 with PDT for GBM treatment using xenograft GBM models implanted in the flank of mice and virally infected with an oHSV1 expressing a membrane-targeted KR variant [3].

In the present study, we generated oHSV1 variants expressing mitochondria-targeted KR. We hypothesized that mitochondrial localization would enhance immunogenic cell death (ICD) more effectively than membrane-localized KR by inducing oxidative stress, releasing damage-associated molecular patterns (DAMPs), and amplifying immune activation [30]. Mitochondria play a central role in ICD by regulating apoptosis, oxidative stress, and inflammation [31, 32]. Unlike membrane-targeted KR, which primarily damages the plasma membrane, mitochondrial KR is expected to robustly activate these pathways. Given that ICD is particularly effective in cancer therapy [33] and that OVs, including oHSV1, rely on ICD to enhance antitumor immunity [34, 35], mitochondrial KR could synergize with oHSV1 by maximizing oxidative stress and immune activation, thus improving viral oncolysis.

To further potentiate the immune response, we also engineered an oHSV1 variant co-expressing KR with immunotherapeutic factors. Specifically, we selected an immune checkpoint inhibitor (anti-PD1 antibody) [36] and interleukin-12 (IL-12), a proinflammatory cytokine known to enhance adaptive immunity in OV therapy. IL-12 stimulates cytotoxic T cells and NK cells, exhibits anti-angiogenic properties, and has demonstrated potent antitumor activity in various cancer models [37]. We evaluated the cytolytic effects of these recombinant oHSV1s *in vitro* using human and murine GBM cell lines and patient-derived cells. Additionally, we assessed their safety and therapeutic efficacy in syngeneic immunocompetent C57BL/6J mice, including animals with GL261-derived brain tumors.

## Results and Discussion

### Oncolytic HSV1s Bearing the mir124 Target Sequences Can Efficiently Infect and Kill Human and Murine GBM Cells While Their Replication Is Impaired in Neurons

To integrate oncolytic virotherapy with PDT, we engineered an oncolytic HSV-1 (oHSV1) expressing the photosensitizer KR protein [27]. Our starting point was an oHSV1 with a genetic backbone similar to the clinically approved T-VEC [38], featuring deletions in the γ34.5 and Us12 viral genes [39, 40]. Unlike T-VEC, our construct lacks the two copies of the human granulocyte-macrophage colony-stimulating factor (GM-CSF) gene. Instead, it harbors a cassette encoding firefly luciferase (FLuc), inserted into the UL55-UL56 intergenic region under the control of the human cytomegalovirus (CMV) immediate early promoter.

To further enhance neuroattenuation beyond the γ34.5 deletion, we introduced target sequences for two forms of mir124 microRNA, which is highly expressed in mature neurons [41]. These sequences were placed downstream of the UL29 gene, encoding an essential viral replication protein [42].

Using oHSV1-mir124 as the parental virus, we then: (i) Replaced FLuc with mitochondria-targeted KR, achieved by fusing the KR-encoding sequence with the cytochrome c oxidase (COX) 8 presequence (oHSV1-KR); (ii) Generated oHSV1-mCherry, where KR was replaced with the mCherry fluorescent protein; (iii) Developed oHSV1-KR-mIL12-aPD1, derived from oHSV1-KR, by inserting a cassette encoding murine interleukin-12 (mIL12) and a single-chain anti-PD1 antibody (aPD1) into one of the γ34.5 loci, under the control of the RFPL4b promoter.

Next, we assessed the infectivity and cytolytic activity of our oHSV1 variants in human U87-MG and LN229 GBM cell lines [43]. Both were highly susceptible to infection, leading to cell death within 72 hours post-infection (**Figure 1A-D**). To further evaluate viral efficacy in a 3D culture model, we infected U87-MG spheroids with oHSV1-mCherry. As shown in **Figure 1E-F**, the virus effectively infected and eliminated GBM cells. These findings were corroborated using patient-derived GBM cells, where oHSV1 induced a marked reduction in cell viability over time (**Figure 2**).

**Figure 1.**
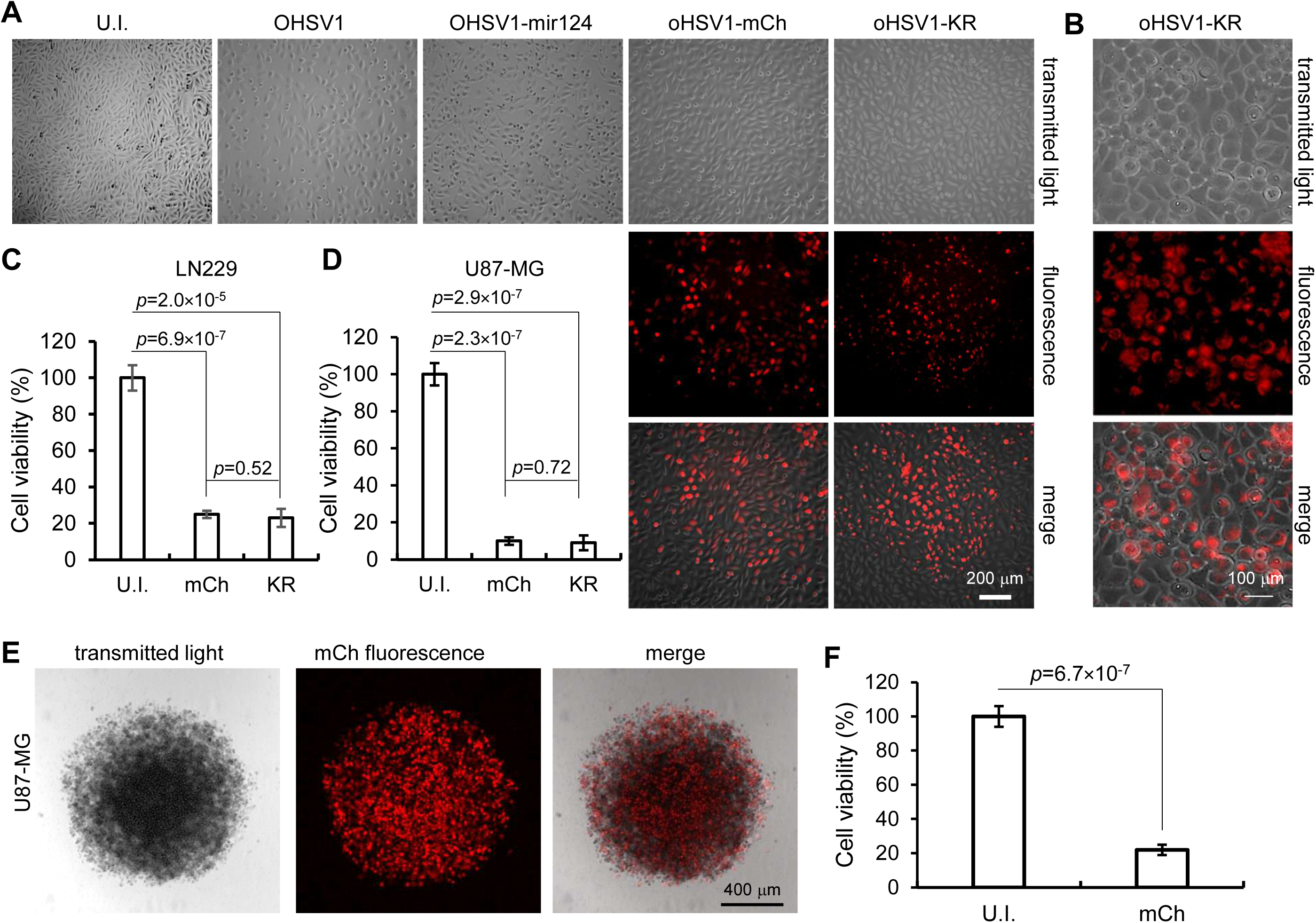
oHSV1 infects and kills human GBM cells. (A, B) LN229 cells infected with different recombinant oHSV1 at MOI=0.1 plaque forming units (PFU) per cell. U.I., uninfected; oHSV1, oncolytic herpes simplex virus type 1; oHSV1-mir124, oHSV1 enriched with UL29/mir124; mCh, mCherry; KR, KillerRed (C, D) Cell viability (trypan blue exclusion assay) at 72 h post infection (p.i.) expressed as % relative to uninfected (U.I.) controls (mean ± s.d., pooled data, n=3 independent experiments). (E) U87-MG tumor spheroid infected with oHSV1-mCh (5’10^5^ PFU, day 7 p.i.). (F) Viability of infected tumor spheroid cells (MTT assay, day 7 p.i.).

**Figure 2.**
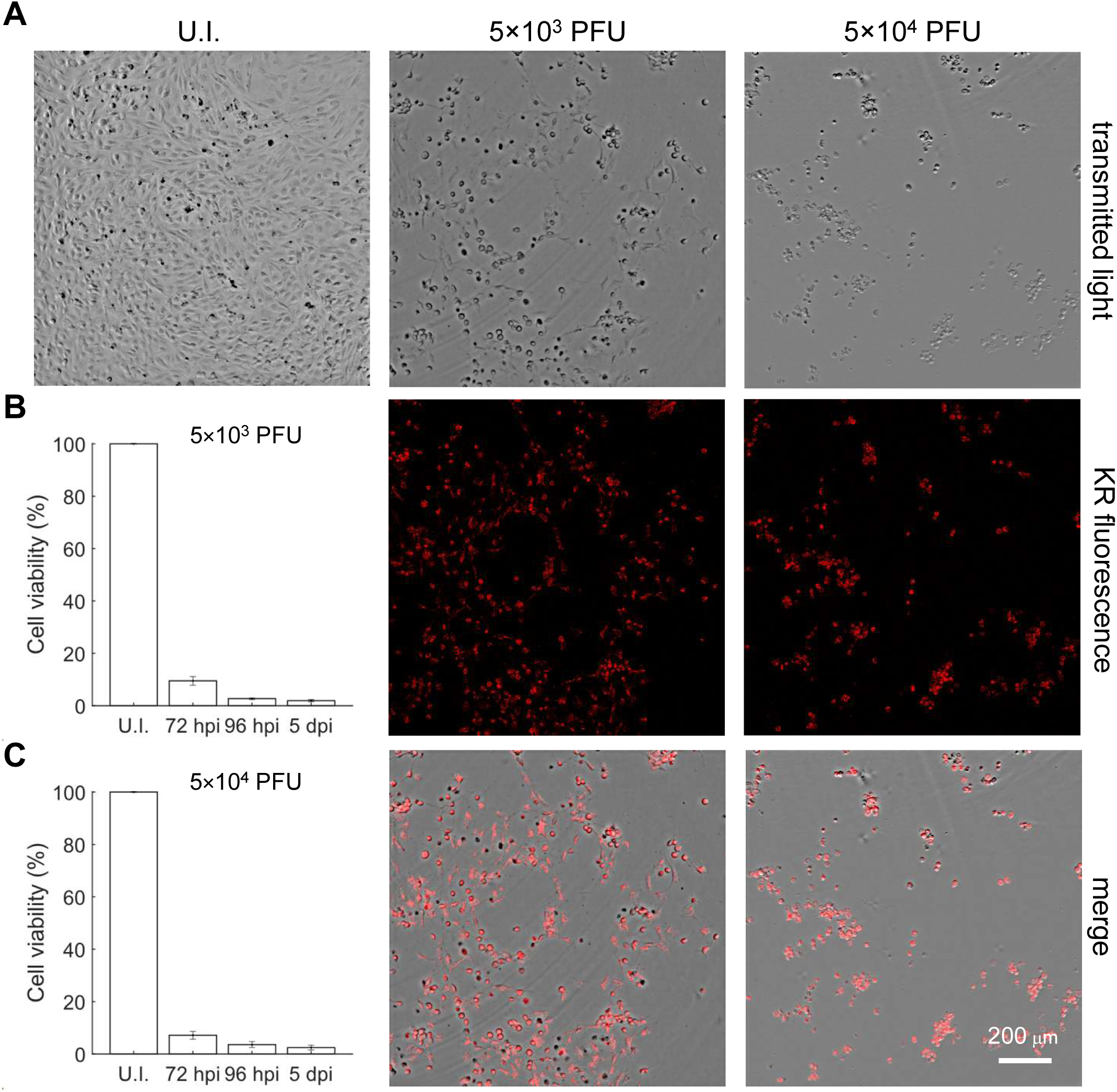
oHSV1-KR infects and kills patient-derived GBM cells. Cells were infected with the indicated PFU of oHSV1-KR. (A) Shown confocal images are representative of the spreading of viral infection (KR fluorescence and merge panels) and cytopathic effect (transmitted light and merge panels). U.I., uninfected cells (control). (B, C) Cell viability was estimated by image segmentation using ImageJ to detect cell contours [57].

To assess the neuroattenuation of our constructs, we compared their replication efficiency to wild-type HSV-1 (strain 17) and oHSV1-Fluc (lacking mir124 target sequences) in neurons. Cells were infected with 10^4^ PFU of each virus, and viral titers were quantified at 48 hours post-infection. Wild-type HSV-1 replicated efficiently in neurons, reaching titers of ∼10⁵ PFU/mL. Recombinant oHSV1 variants displayed significantly reduced replication, with oHSV1-Fluc reaching ∼10³ PFU/mL and oHSV1-mCherry showing an even lower titer (∼10² PFU/mL). Expression of both KR and mCherry was minimal in neurons (**Figure 3**), with only 1% and 3% of positive cells, respectively, confirming efficient neuroattenuation via mir124 targeting.

**Figure 3.**
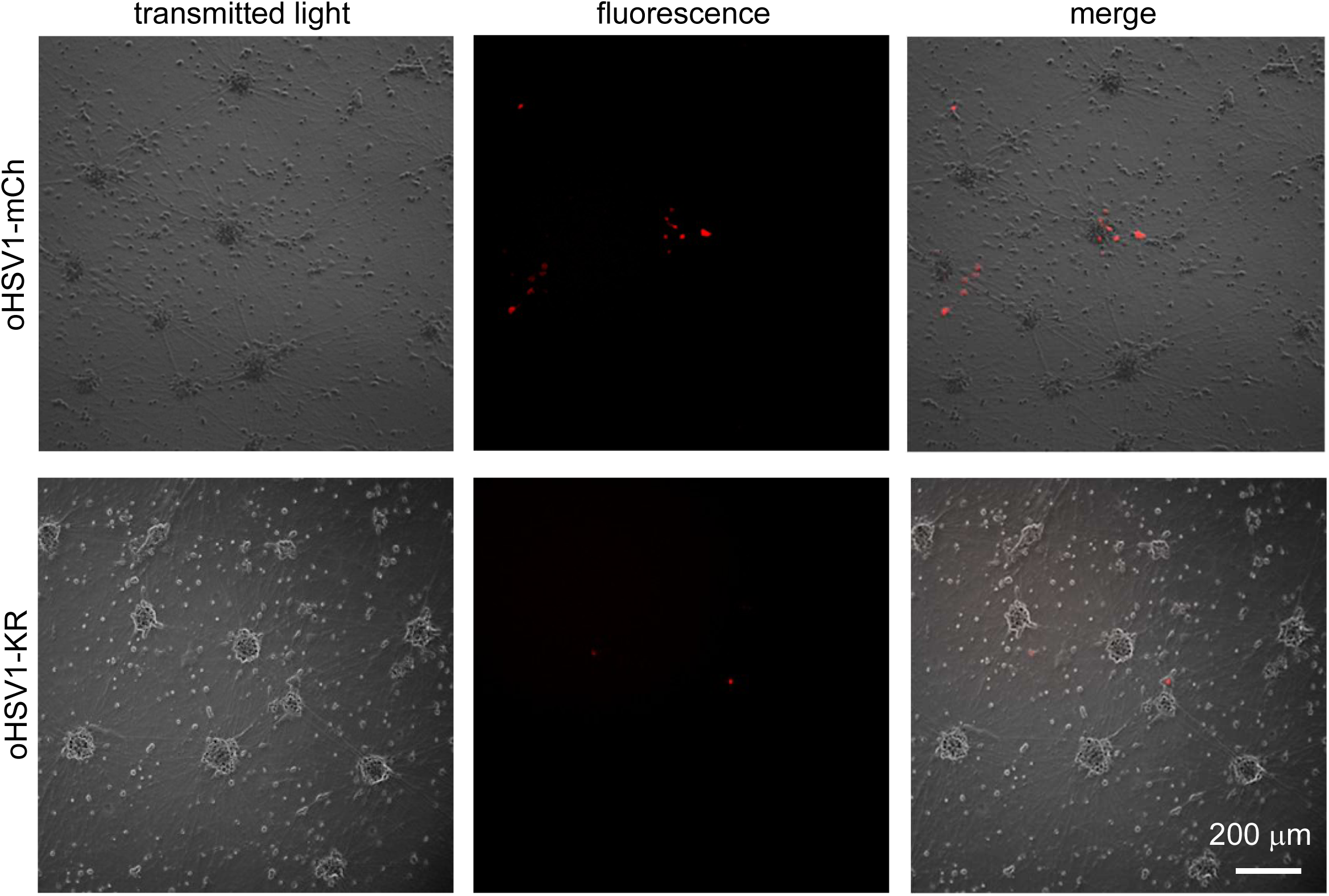
oHSV1-KR is attenuated in neurons. Neurons derived from pluripotent stem cells were infected with the same amount (10^3^ PFU) of the indicated oHSV1s. Shown are representative images at 48 hours p.i.. The low percentage of red fluorescent cells is due to virus neuroattenuation; compare to Figure 1A and 1B.

### Viral Spread and Cytotoxicity in Murine GBM Models (GL261)

Recombinant viruses efficiently infected and spread in GL261 murine GBM cells, both in 2D and 3D cultures (tumor spheroids), inducing significant tumor cell death (**Figure 4A–C**). All tested GBM cell lines and primary cells showed robust KR and mCherry expression. KR expression in GL261 cells was further quantified using microplate reader analysis of fluorescence emission, demonstrating a clear correlation between viral MOI and reporter protein expression (**Figure 4D**). To verify the functional expression of mIL12 and aPD1, we also performed Real-Time PCR on cDNA extracted from oHSV1-KR-mIL12-aPD1-infected GBM cells (CT of 27 and 32 for mIL12 in LN229 and GL261, respectively; CT of 30 and 34 for aPD1 in LN229 and GL261, respectively). Furthermore, mIL12 protein secretion was quantified by ELISA in the supernatant of highly susceptible Vero cells, revealing a concentration of ∼800 ng/mL, which is indicative of efficient protein production and release upon infection.

**Figure 4.**
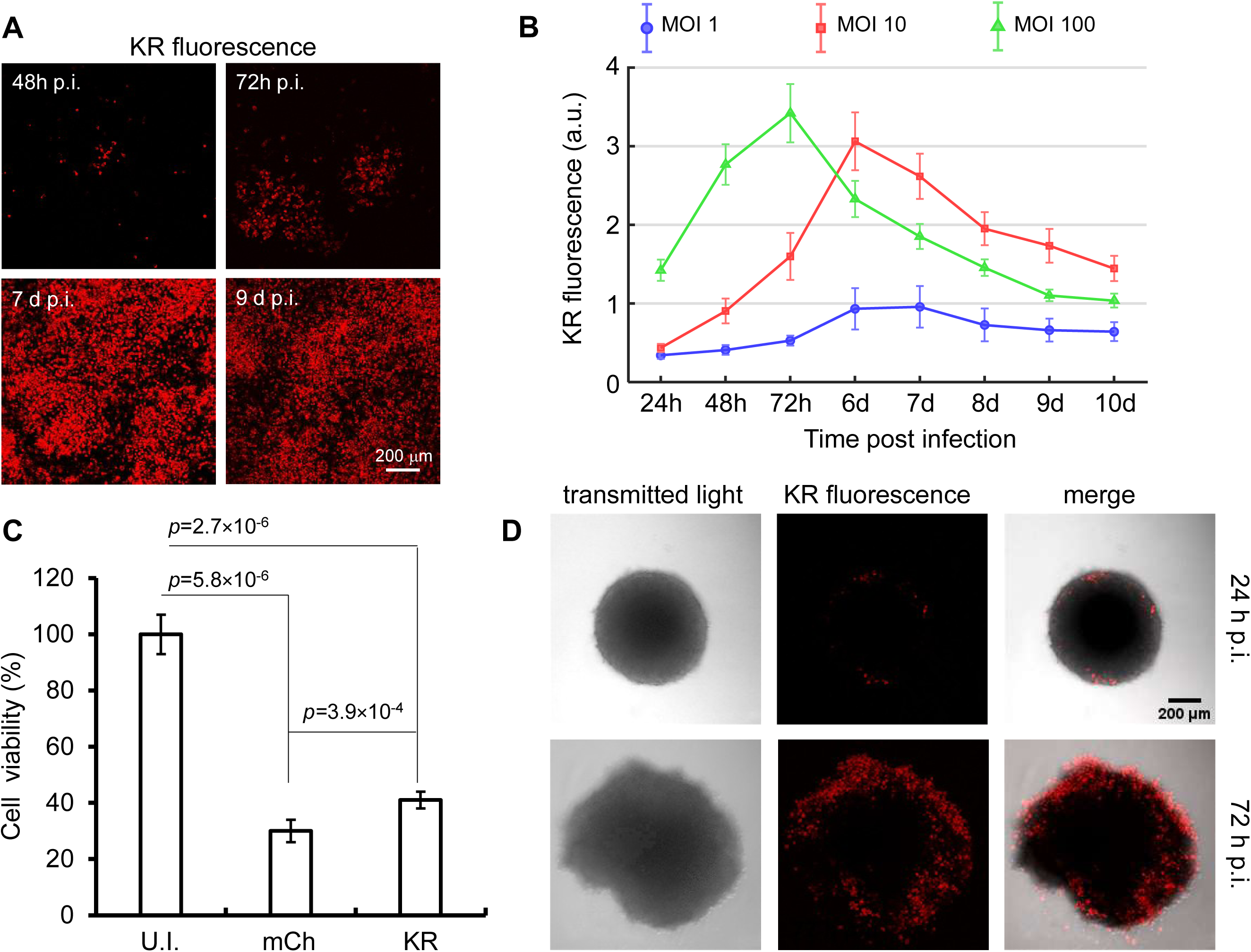
oHSV1 infects, spreads in, and kills mouse GL261 cells. (A) Representative confocal fluorescence images of GL261 cells infected with oHSV1-KR (MOI =10 PFU/cells) and observed at the indicated times post infection (p.i.). (B) Plate reader quantification (96-well plates) of KR fluorescence level in oHSV1-KR infected GL261 cells infected (MOI=1, 10, 100 PFU/cell). The decreases of KR fluorescence after MOI-dependent peak are due to the cytopathic effect of the virus. (C) Viability of GL261 cells infected (MOI=10 PFU/cells) with oHSV1-mCherry (mCh) or oHSV1-KR (KR); trypan blue exclusion assay performed 10 days (d) p.i. and quantified as % of the uninfected controls (U.I.). (D) Confocal microscopy images of GL261 spheroids infected with oHSV1-KR (6×10^5^ PFU).

### Monocytes as Carriers for Systemic oHSV1 Delivery to GBM Cells

With the aim of developing a feasible approach for the systemic delivery of oHSV1 *in vivo*, we recently proposed the use of autologous monocytes [39]. We showed that oHSV1-loaded primary human monocytes migrated *in vitro* towards epithelial cancer cells of different origin. Moreover, human monocytic leukemia cells selectively delivered oHSV1 to human head-and-neck xenograft tumors grown on the chorioallantoic membrane (CAM) of fertilized chicken eggs after intravascular injection [39].

Starting from these data monocytes isolated from buffy coats of healthy donors were infected with oHSV1-mCherry at the MOI of 5 PFU/cells. Migration assays revealed that oHSV1-infected monocytes efficiently migrate towards cell media conditioned by human GBM cells (**Figure 5A**), Furthermore, when infected monocytes were cultured in the same media the amount of infectious particles released in the supernatants increased over time, as quantified by plaque assays (**Figure 5B**). Finally, monocytes loaded with oHSV1-mCherry successfully transmitted the virus to U87-MG spheroids, leading to efficient tumor cell killing (**Figure 5C-D**).

**Figure 5.**
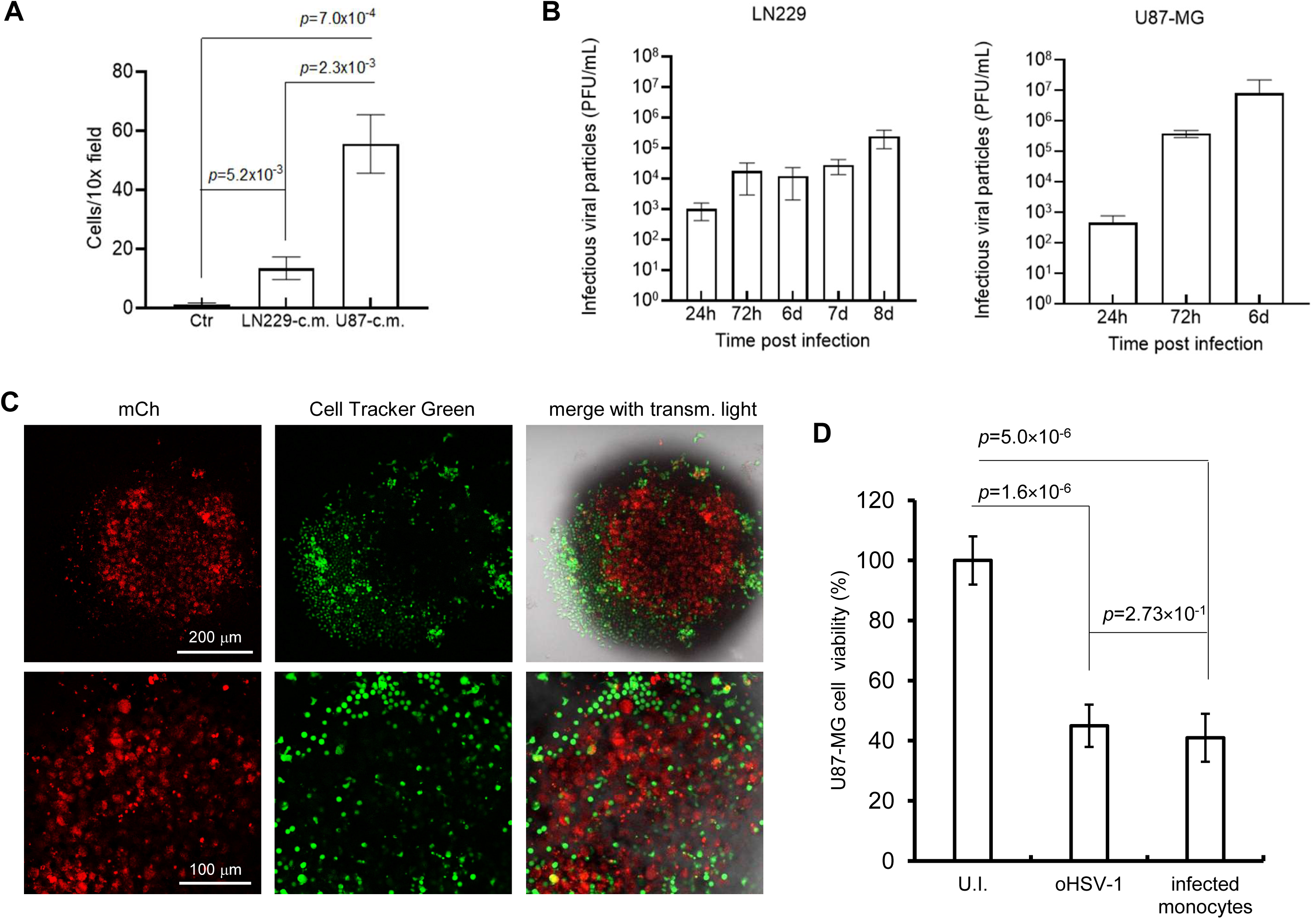
Primary human monocytes loaded with oHSV1-mCherry transmit viral infection to human GBM cells, reducing their viability. (A) Primary human monocytes infected with oHSV1-mCherry at the MOI of 5 PFU/cell were allowed to migrate towards serum-free medium conditioned by human GBM cells, as indicated, through a 5.0 µm-pore filter for 3 h. Cells were counted from at least three independent fields of view in the lower chamber. (B) Infected monocytes were cultured at a 1:1 ratio with confluent U87-MG or LN229 cells, as indicated. Supernatants were harvested at the indicated time points and titrated via plaque assay on Vero cells. (C-D) U87-MG spheroids were treated with human primary monocytes infected with oHSV1-mir124-mCherry (MOI = 3 PFU/cell) and labeled with CellTracker™ Green (5×10^3^ cells/well). Confocal microscopy analysis was performed at 7 days post treatment (C) and cell viability was assessed by MTT assay (D). The bar graph displays the mean values obtained from three replicates and the corresponding standard deviation. Infection with free oHSV1 produced a comparable reduction of cell viability.

To assess whether murine monocytes could also serve as viral carriers, we performed co-culture experiments with GL261 murine GBM cells at a 1:1 monocyte-to-tumor cell ratio. Over several days of confocal microscopy observation, we observed a progressive increase in the number of red fluorescent cells, confirming the successful transfer of oHSV1-mCherry to GL261 cells (**Figure 6**). Importantly, cell viability assays at 13 days post-treatment revealed complete eradication of tumor cells, with no viable cells detected, while roughly 1×10^6^ alive cells were found in the negative control (uninfected GL261).

**Figure 6.**
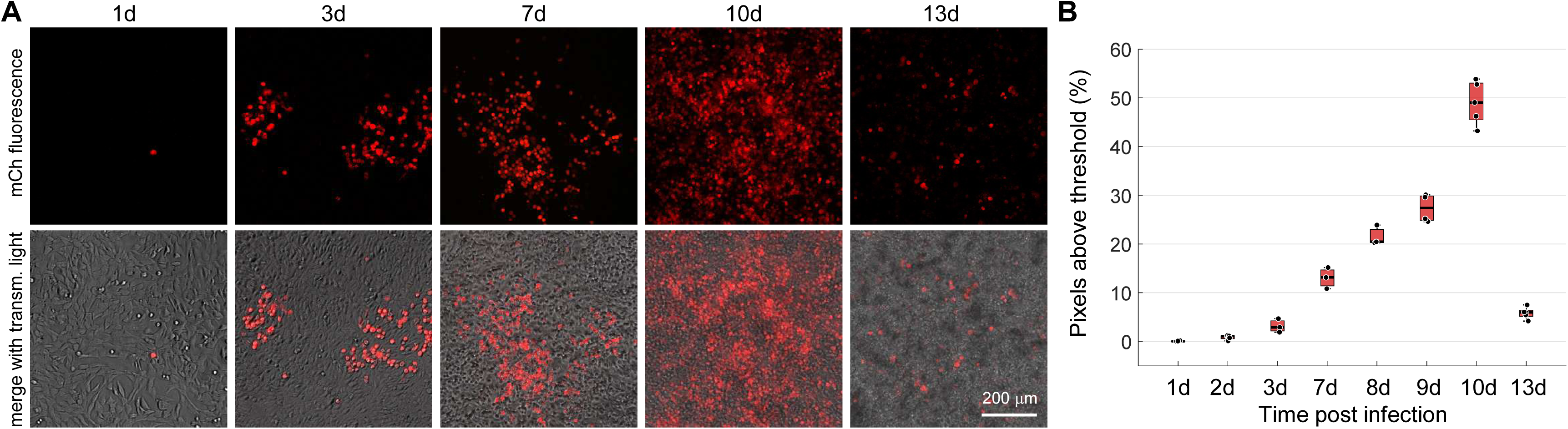
Murine monocytes loaded with oHSV1-mCherry transmit viral infection to mouse Gl261 cells. (A) Infected monocytes (MOI = 3 PFU/cell) were seeded over GL261 cells (3×10^5^/well) and expression of mCherry (mCh) in infected cells was tracked by fluorescence microscopy at the indicated days (d) p.i.. (B) Dot-box plots showing % of image pixels with mCh fluorescence above a fixed arbitrary threshold vs. time p.i.. For each time point, n ≥ 3 independent fields of view were analyzed using Fiji/ImageJ and the number of pixels above a fixed threshold was normalized to the total. The sharp drop at 13d is due to the cytopathic effect of the virus.

Overall, these findings strongly support the feasibility of monocytes as systemic carriers for oHSV1 delivery *in vivo*. This approach could overcome current limitations associated with intracranial injection, enabling less invasive and potentially repeatable administration of oncolytic virotherapy for GBM treatment.

### Mitochondria-Targeted KR Reduces GL261 Cell Viability Only Upon Photoactivation

Next, we evaluated the impact of mitochondria-targeted KR on murine GBM (GL261) cell viability, both before and after photoactivation. To achieve this, we constructed lentiviral vectors encoding either KR or the EGFP derived glutamate sensor iGluSnFr under the transcriptional control of the same promoter RFPL4b. These constructs were used to generate VSV-G-pseudotyped recombinant lentiviral particles (RLPs), which were subsequently used to transduce GL261 cells. Stably transduced cells were then selected using puromycin, expressed by the vectors. We assessed the growth kinetics of GL261 cells stably expressing KR (GL261-KR) comparing them to control GL261 cells, i.e. un-transduced GL261 cells (GL261-WT) and GL261 cells stably expressing iGluSnFr (GL261-iGlu). Proliferation rates (**Figure 7A**) remained comparable across all cell lines, confirming that KR expression alone does not affect cell viability. In contrast, signs of apoptosis appeared upon confocal laser irradiation (75 min, 561 nm, 2.5 mW at the focal spot) of GL261-KR cells **(Figure 7B**). Irradiating GL261 cells infected with oHSV1-KR at an MOI of 10 PFU/cell significantly reduced cell viability compared to infected cells that were not subjected to irradiation (**Figure 7D**).

**Figure 7.**
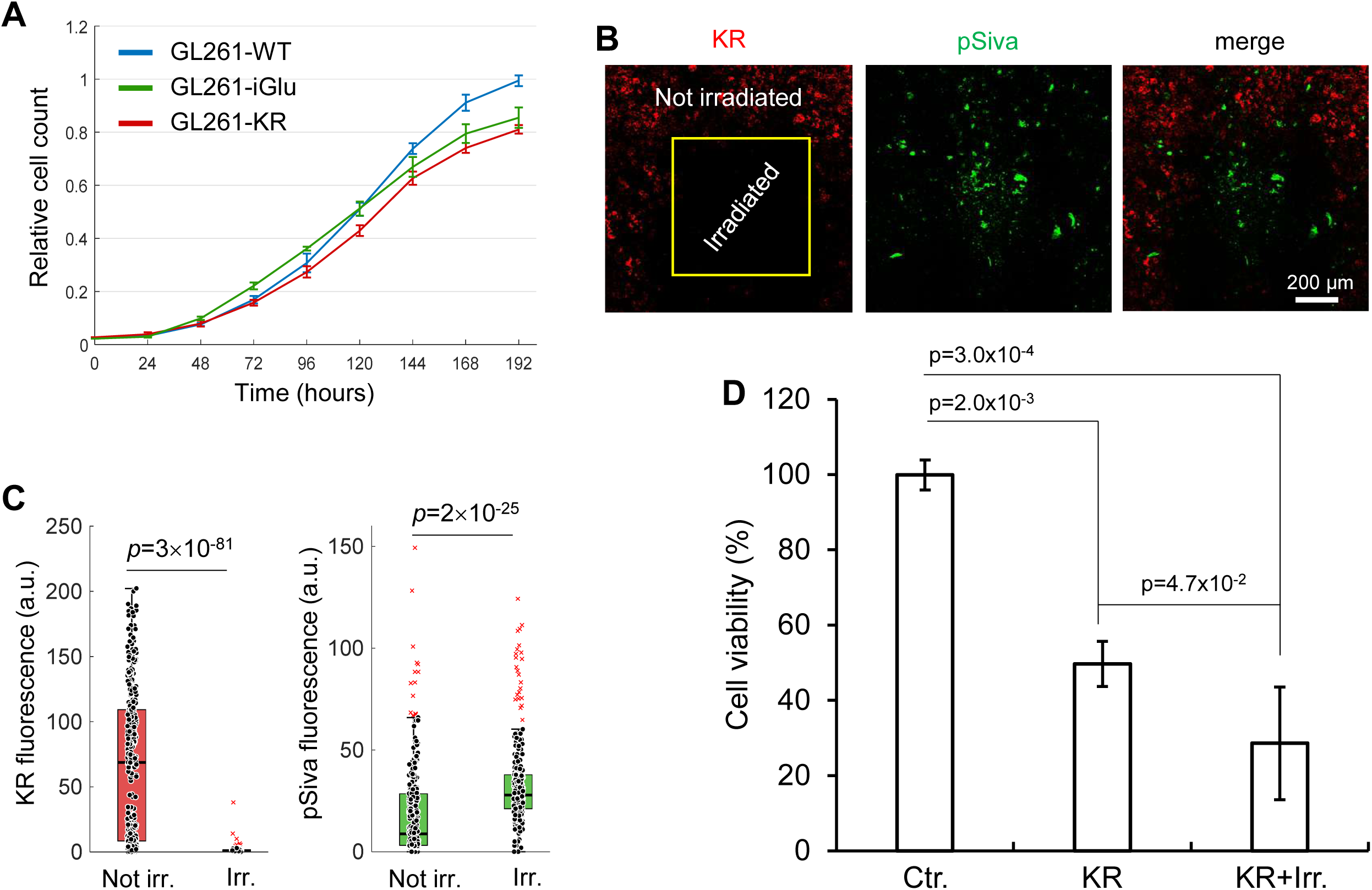
Growth of GL261-derived stable cell pools and effect of KR photoactivation in 2D cultures. (A) Growth curves of GL261-WT, GL261-KR and GL261-iGluSnFR cells in 2D cultures. (B) Irradiation of GL261-KR cells (561 nm, 2.5 mW at focal point, 75 min). (C) Dot-box plots of KR and pSiva fluorescence intensity in irradiated cells (Irr., n = 284) and not irradiated cells (n = 350) 3 h post irradiation. (D) Viability of GL261 cells infected with oHSV1-KR (KR) (MOI=10 PFU/cell) irradiated or not with a 565 nm LED for 30 min (120 mW/cm^2^) and quantified by trypan blue exclusion assay 48 h p.i.; Ctr.: not infected, not irradiated controls.

We also established three-dimensional (3D) cultures of GL261 cells and neurons/astrocytes using Alvetex scaffolds (**Figure 8A-G**). GL261-KR spheroids seeded over astrocyte-populated scaffolds were used to set the conditions for the photoactivation of the KR protein targeted to the mitochondria in tumor cells *in vivo.* Spheroids were loaded with calcein AM (4 mM, 5 min), bathed in a CO_2_-idependent culture medium supplemented with DAPI (5 mM) and maintained at 37 °C. KR proteins were photoactivated with 588 nm laser light delivered by a tapered fiber optics with a 10 micron tip placed in the proximity of the spheroid (90 mW at fiber output). Real-time confocal fluorescence imaging revealed loss of calcein paralleled by uptake of DAPI, both indicative of progressive reduction of plasma membrane integrity due to KR photoactivation (**Figure 8H-O**).

**Figure 8.**
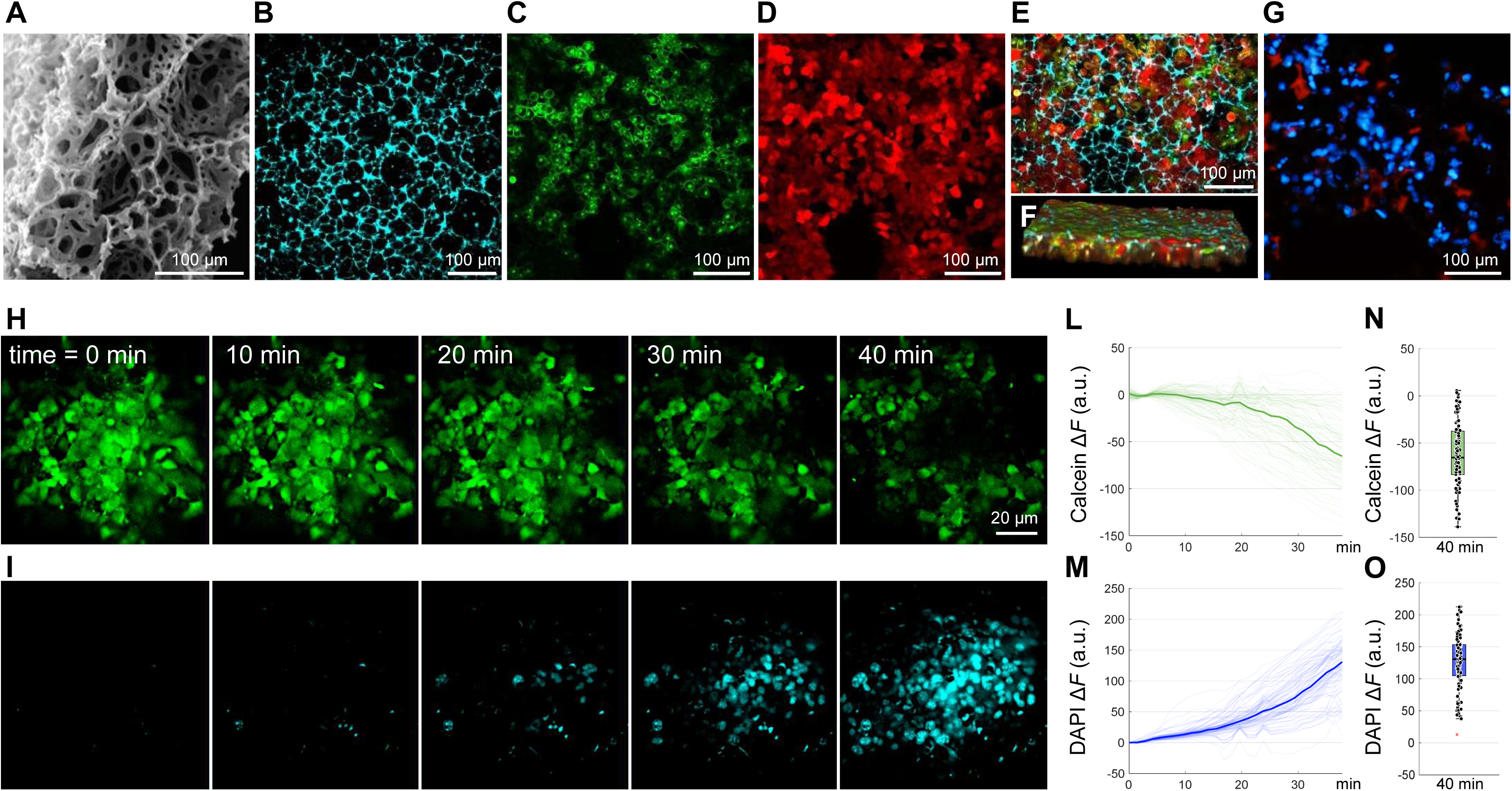
Photoactivation of GL261-KR tumor spheroids seeded over Alvetex scaffold populated with astrocyte reduces plasma membrane integrity. (A) Scanning electron microscope image of 200 μm thick Alvetex Scaffold disc characterized by voids with dimensions of 42 μm in diameter and interconnects of 13 μm in diameter (source: https://www.reprocell.com/alvetex). (B) Cyan-colored structure is the Alvetex Scaffold visualized by second-harmonic generation signals. (C-F) Two-photon fluorescence images of GL261 cells grown for 10 days in Alvetex Scaffold. Cells are marked in green (C) and red (D). 3D views of the scaffold invaded by the cells in (E) the focal plane (xy) and (F) orthogonal planes (xz). (G) Cryosection of an Alvetex scaffold populated with neurons and astrocytes. Nuclei were stained with DAPI, and cells were marked with a red vital dye. (H-I) Confocal live imaging frames of a GL261 tumor spheroid pre-loaded with Calcein AM (H) and bathed in a CO_2_-idependent culture medium supplemented with 5 mM DAPI (I) while KR was photoactivated with 588 nm laser light delivered by a tapered fiber optics with a 10 micron tip placed in the proximity of the spheroid. (L, M) Thin lines: fluorescence emission variation (Δ*F*) of Calcein (L) and DAPI (M) vs. irradiation time in n = 105 cells of the spheroid; thick lines: population medians. (N, O) Dot-box plot quantification of Δ*F* at the end of the 40 min irradiation period.

### Oncolytic HSV1-mCherry Can Be Safely Delivered *in Vivo*

To evaluate the safety of oHSV1-mCherry *in vivo*, we conducted a proof-of-principle study in which 1.8×10⁵ PFU of the virus were injected directly into the striatum of healthy adult male C57BL/6J mice. Brains were collected at 7 days (n = 3) and 14 days (n = 3) post-injection.

Throughout the observation period, mice were monitored daily, and no significant changes in body weight were detected (**Figure 9A)**. However, two mice exhibited mild neurological symptoms: one displayed tremors and lethargy on day 6 (16.6%), while another exhibited lethargy on day 8 (16.6%). To assess viral diffusion, we analyzed native mCherry fluorescence and confirmed infection through immunofluorescence staining using anti-mCherry antibodies (**Figure 10A, B**). Robust mCherry expression was observed at 7 days post-injection, which remained high at 14 days, demonstrating that oHSV1-mCherry spreads around the injection site and infects neurons, including their cell bodies, dendrites, and axon terminals. These findings are consistent with previous reports [44].

**Figure 9.**
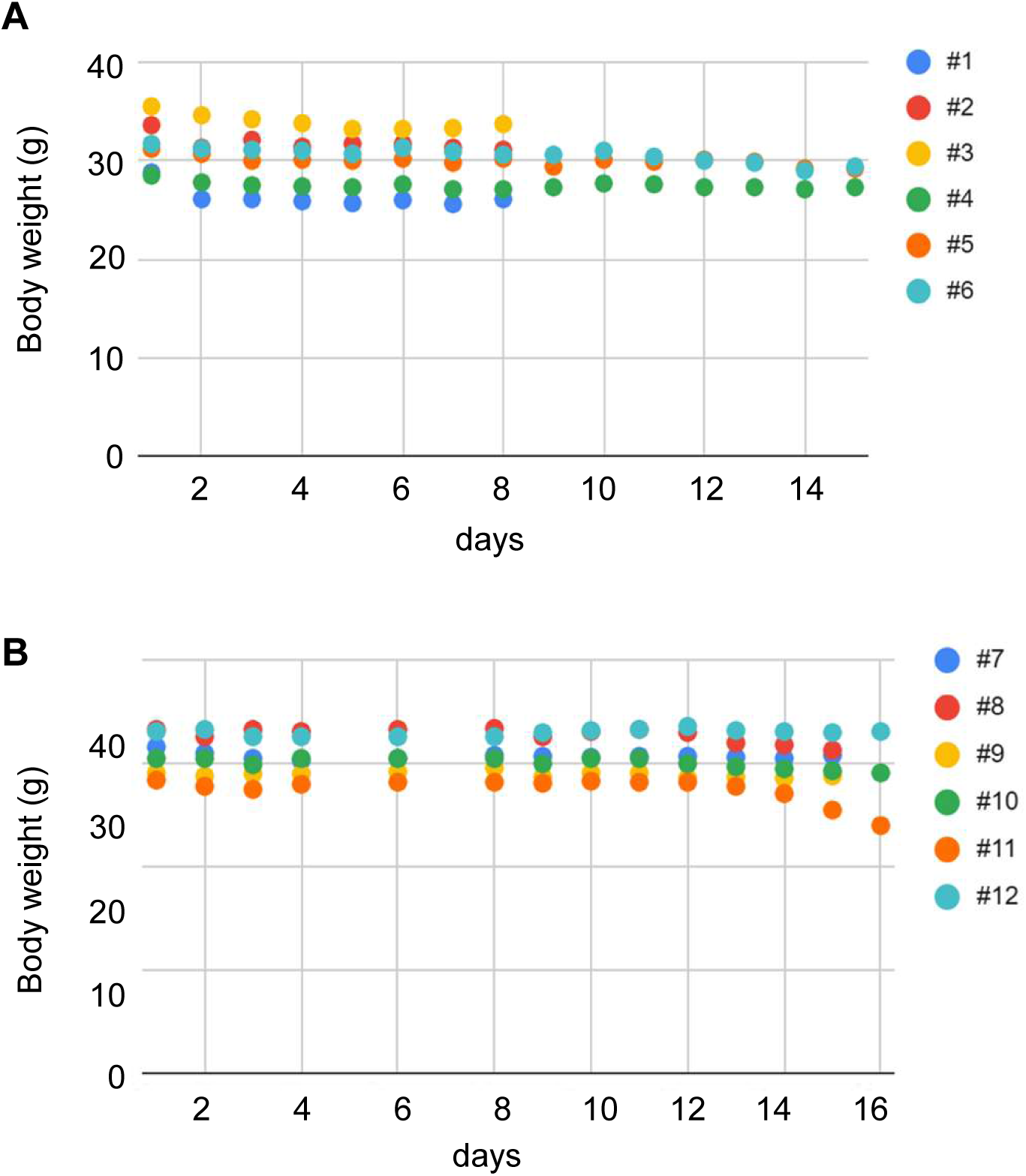
Body weight curves of mice treated with oHSV1-mCherry alone or with infected-monocytes. (A) Body weight of healthy adult male C57BL/6J mice after injection with oHSV1-mCherry measured for 7 days (blue, red and yellow dots) or 14 days (green, orange and light blue dots). (B) Body weight of mice bearing intracranial GL261 tumors after injection with oHSV1-mCherry alone (blue, red and yellow dots) or with infected monocytes (green, orange and light blue dots). Numbers besides colored dots are mouse ID numbers.

**Figure 10.**
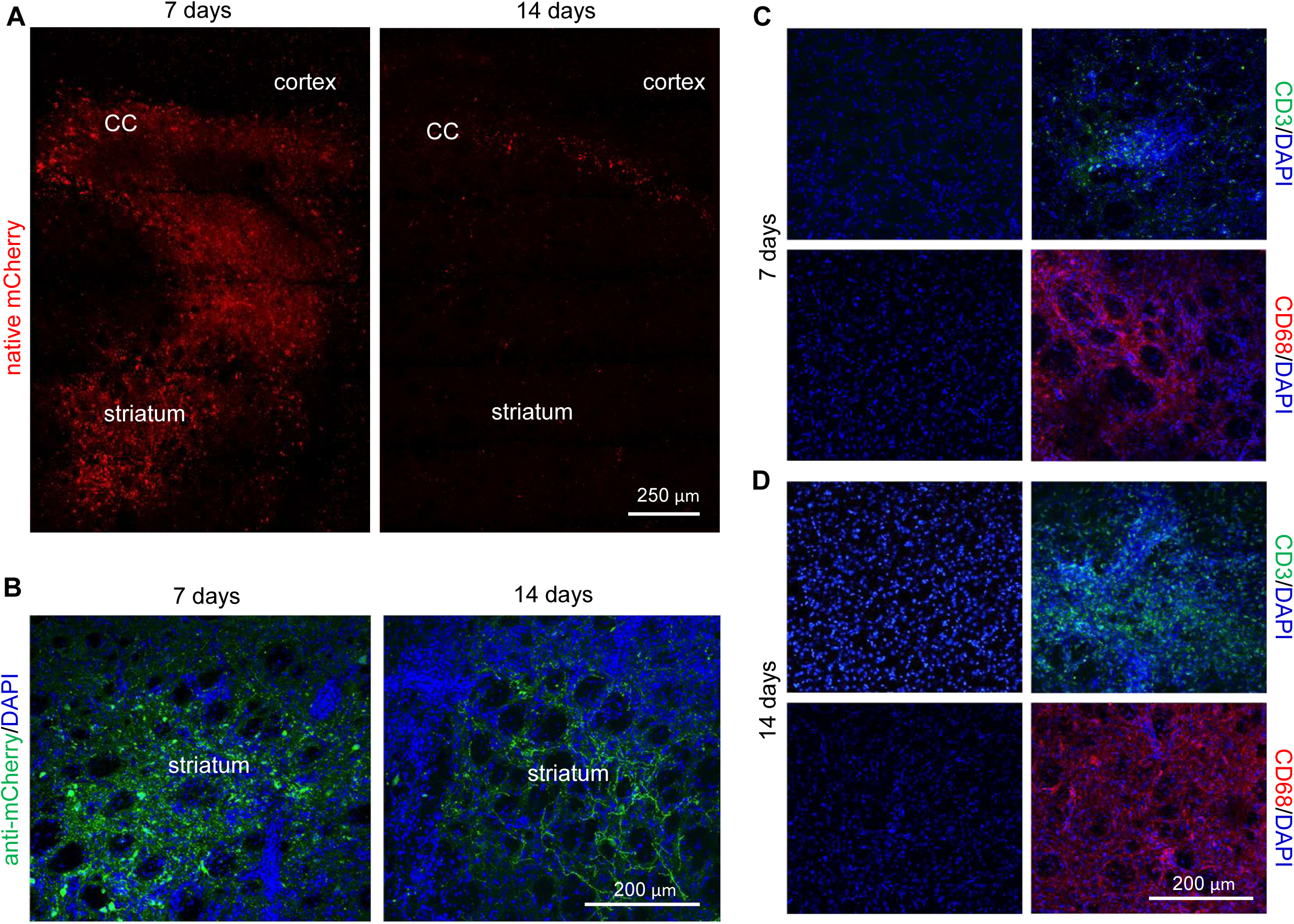
oHSV1-mCherry expression and CD3+ and CD68+ cells infiltration in healthy mouse brains after oHSV1-mCherry injection. (A, B) Representative images of coronal sections from animals sacrificed 7 days or 14 days after oHSV1-mCherry injection in the right striatum (see Materials and Methods for details). Native mCherry fluorescence imaged using multiphoton microscopy (A). Immunofluorescence images of striatum using an anti mCherry antibody (green). Nuclei are stained with DAPI (blue; B). (C, D) Representative immunofluorescence images of coronal sections from animals sacrificed 7 days (C) or 14 days (D) after oHSV1-mCherry injection in the right striatum using CD3 (green) or CD68 (red) in the non-injected (left panels) or injected (right panels). Nuclei are stained with DAPI (blue). CC: corpus callosum.

Additionally, we assessed immune cell infiltration using CD3 (T cells) and CD68 (microglia/macrophages) immunostaining (**Figure 10C, D**). Both markers were present at day 7 post-injection, with an increase in CD3+ and CD68+ cell infiltration at day 14, indicating a progressive immune response. Overall, these results suggest that the virus diffused around the injection site, inducing infiltration of immune cells.

### Intratumoral Administration of oHSV1-mCherry Alone or Carried by Monocytes Induces Tumor Necrosis, CD3/CD68+ Cell Infiltration and Neuroinflammation

To further evaluate therapeutic effects, we injected oHSV1-mCherry alone (n = 3) or oHSV1-infected monocytes (n = 3) into the right striatum of mice 7 days after GL261 tumor cell implantation (6×10^4^ cells/mouse). Mice were monitored daily, and only one mouse (16.6%) exhibited transient weight loss and lethargy within 2 hours post-injection of infected monocytes (**Figure 9B**). Brains were collected on day 21 post-tumor injection and histological analysis revealed that tumors treated with oHSV1-mCherry alone exhibited large necrotic areas in the tumor core, accompanied by significantly increased infiltration of CD3+ T cells compared to tumors treated with infected monocytes (**Figure 11A-C**). Quantification of CD68 fluorescence intensity was comparable between the two conditions (**Figure 11B, D**). Neuroinflammatory responses were analyzed by Gfap (astrocyte reactivity) and Iba1 (microglia-macrophage activation) immunostaining. Neuroinflammation was most prominent within the tumor microenvironment but also extended throughout the brain, and significant differences were found for Gfap in the peritumoral area and Iba1 in the tumor mass for oHSV1-mCherry alone compared to oHSV1-infected monocytes (**Figure 11E-G)**. Overall, these results suggest that virus alone or carried by monocytes elicit distinct tumor-immune microenvironmental response.

**Figure 11.**
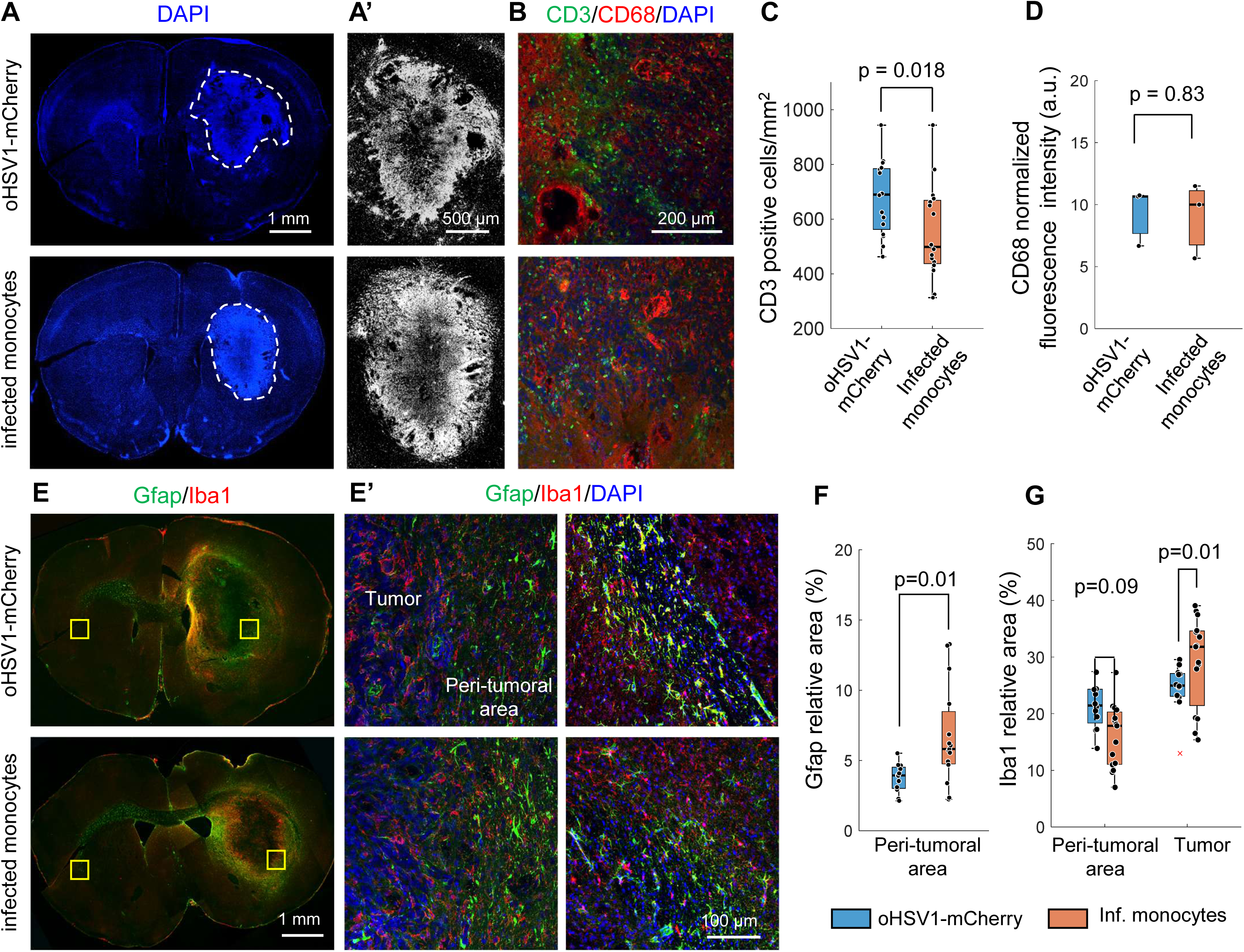
Tumor growth, CD3+ and CD68+ cells infiltration and neuroinflammation in C57BL/6J mice implanted with GL261 cells treated with oHSV1-mCherry alone or with infected-monocytes. Representative DAPI staining (A, blue; enlarged images of tumor areas, A’, grey) of coronal sections from C57BL/6J mice implanted with GL261 cells treated with oHSV1-mCherry alone (left) or with infected-monocytes (right). (B). Representative co-immunofluorescence images of CD3 (green) or CD68 (red) of tumor areas from coronal sections from C57BL/6J mice implanted with GL261 cells treated with oHSV1-mCherry alone (left) or with infected-monocytes (right). Nuclei are stained with DAPI (blue). Dot-box plot quantification of CD3+ T cells (C) and CD68 fluorescence intensity (D) from n=3 mice for each experimental group. (E) Representative Gfap (green) and Iba1 (red) co-immunostaining of coronal sections from C57BL/6J mice implanted with GL261 cells treated with oHSV1-mCherry alone (left) or with infected-monocytes (right). E’ are enlarged images from E of peri-tumoral (upper panels) or contralateral uninjected areas (lower panels). Nuclei are stained with DAPI (blue). Dot-box plot quantification of Gfap (F) and Iba1 relative areas (D) from n=3 mice for each experimental group.

## Conclusions

In this study, we developed and characterized novel oncolytic HSV-1 (oHSV1) variants engineered for enhanced safety, tumor selectivity, and combinatorial therapeutic efficacy against glioblastoma (GBM). Our findings demonstrate that oHSV1-mCherry is highly neuroattenuated, effectively infecting and killing human and murine GBM cells while exhibiting limited replication in neurons, a crucial feature for ensuring safety in potential clinical applications. By incorporating mir124 target sequences, we achieved a substantial reduction in viral replication in neuronal cells, further enhancing the neuroattenuated profile of our virus.

To overcome the limitations of intracranial oHSV1 administration, we explored monocytes as systemic carriers for viral delivery. Our results confirm that oHSV1-infected monocytes successfully transfer the virus to GBM cells *in vitro*, leading to potent tumor cell killing, even in 3D culture models. Importantly, systemic administration of oHSV1-infected monocytes in mice via intravenous (i.v.) or intraperitoneal (i.p.) injection was well tolerated, with no significant signs of distress or off-target viral dissemination. These results highlight the potential of monocyte-based systemic delivery strategies to enable repeated, minimally invasive oHSV1 administration.

In addition to viral-mediated cytotoxicity, we introduced mitochondria-targeted KillerRed (KR) to further enhance photodynamic therapy (PDT)-induced tumor cell death. Our *in vitro* studies demonstrate that KR expression does not affect GL261 cell viability in the absence of light activation, ensuring the controllability of PDT. Moreover, KR-expressing oHSV1 variants, combined with immunostimulatory factors such as IL-12 and anti-PD1, offer an innovative approach to enhance antitumor immune responses.

Finally, in preclinical *in vivo* models, intracranial injection of oHSV1-mCherry into GBM-bearing mice resulted in large necrotic areas within the tumor core and increased CD3+ immune cell infiltration, indicating a strong immune activation. In contrast, tumors treated with monocyte-delivered oHSV1 exhibited a different morphology, suggesting a distinct tumor-immune microenvironmental response.

Overall, our findings establish a strong preclinical foundation for the use of engineered oHSV1 as a multimodal therapeutic platform for GBM. By integrating oncolytic virotherapy, photodynamic therapy, and immune modulation, we provide a rationale for further studies exploring optimized delivery strategies and combination therapies to maximize efficacy. Future investigations will focus on further improving systemic delivery, refining light-based activation of KR *in vivo*, and evaluating long-term antitumor immunity, with the ultimate goal of translating this approach to clinical applications.

## Materials and Methods

### Cells

#### Patient-derived GBM Cells and Ethics

Our research complies with all relevant ethical regulations and guidelines. This study was conducted under protocols approved by the IRBs and IACUCs of the Padova University Hospital (2462P). All patients were provided with written informed consent for all clinical information, treatments, and prospective biopsy acquisition. All tissues were acquired following the tenets of the Declaration of Helsinki.

Cells were isolated from GBM tumors at surgery and cultured as previously described [45]. Briefly, GBM samples were enzymatically and mechanically dissociated into single cell suspensions. Cells were then placed on fibronectin-coated plates and grown as monolayers in DMEM/F12 (Biowest, Nuaillé, France) supplemented with 10% BIT9500 (Stem Cell Technologies, Vancouver, Canada), 20ng/ml basic Fibroblast Growth Factor (bFGF) and 20ng/ml Epidermal Growth Factor (EGF; both from Cell Guidance Systems Ltd, Cambridge, UK). GBM cells were maintained in an atmosphere of 2% oxygen, 5% carbon dioxide and balanced nitrogen in a H35 hypoxic cabinet (Don Whitley Scientific Ltd, Shipley, UK) to better resemble the hypoxic conditions of GBM microenvironment [45, 46].

#### Cell Lines

The following cell lines were used in this study and maintained under standard culture conditions. LN-229 (ATCC, CRL-2611), U87-MG (ATCC, HTB-14™), Vero CCL81 (ATCC, CCL-81™), and HEK-293T (ATCC, CRL-11268) cells were cultured in Dulbecco’s Modified Eagle Medium (DMEM; Gibco, Thermo Fisher Scientific, Monza, Italy) supplemented with 10% (v/v) fetal bovine serum (FBS; Gibco, Termo Fisher Scientific) and 1% (v/v) penicillin-streptomycin (Pen-Strep; Gibco, Thermo Fischer Scientific).

GL-261 (DMSZ, ACC 802) cells were cultured in DMEM high glucose, pyruvate (Gibco, Thermo Fisher Scientific) supplemented with 10% (v/v) FBS (Gibco, Thermo Fisher Scientific), 1% (v/v) Pen-Strep (Gibco, Thermo Fischer Scientific), and 1% (v/v) L-glutamine (Gibco, Thermo Fisher Scientific).

#### 3D Spheroid Cultures

U87-MG and GL261 spheroids were generated as previously described [47]. Briefly, 2×10³ cells per well were seeded in 200 µL of culture medium in Ultra-Low Attachment (ULA) 96-well round-bottom plates (Corning®, New York, NY, USA) and centrifuged at 1000 rpm for 10 min to facilitate spheroid formation.

#### Astrocyte and Neuronal Cultures

Immortalized human astrocytes (HA) (InnoProt, Derio (Bizkaia) Spain, P10251-IM) were maintained in astrocyte medium (InnoProt, P60101) on poly-L-lysine (PLL; InnoProt)-coated flasks. H9 iNe pluripotent stem cells were cultured in mTeSR Plus medium (STEMCELL Techologies, Meda MB, Italy) on Geltrex LEDVE-Free (Thermo Fisher Scientific)-coated multiwell plates. Neuronal differentiation was performed as previously described [48]. Three-dimensional (3D) cultures were established using Alvetex scaffolds following the Alvetex Scaffold: Quick Start Protocol (available at: <u>Reprocell Protocol</u>). Two-photon and second-harmonic generation images of cell-populated scaffolds were acquired with a custom-made multiphoton system previously described in [49].

#### Mycoplasma Testing

To ensure the absence of mycoplasma contamination, cell lines were regularly tested using end-point PCR with AmpliTaq Gold DNA polymerase (Applied Biosystems, Thermo Fisher Scientific). The following primers were used:

- Forward primer: 5’-GGGAGCAAACAGGATTAGATACCCT-3’
- Reverse primer: 5’-CATGTCTCACTCTGTTAACCTC-3’

### Lentiviral Vectors and Generation of Stably Transduced GBM Cells

Third-generation lentiviral vectors (LVs) encoding RFPL4b/KR or RFPL4b/iGluSnFR were obtained from VectorBuilder (Neu-Isenburg, Germany). Both constructs included a puromycin resistance gene for selection of transduced cells.

#### Lentiviral Vector Production

Vesicular stomatitis virus (VSV)-G pseudotyped lentiviral stocks were produced by calcium phosphate transfection of HEK-293T cells, as previously described [50]. Briefly, 12×10⁶ HEK-293T cells were seeded in T-150 flasks 24 hours before transfection and then co-transfected with the following plasmids:

- 20 µg of the gene transfer vector (LV-RFPL4b/KR or LV-RFPL4b/iGlu)
- 10 µg of pMDLg/pRRE (#54)
- 10 µg of pRSV-REV
- 10 µg of pMD2.VSV.G

All plasmids were kindly provided by L. Naldini. Culture supernatants were collected 48 hours post-transfection, filtered through a 0.45-µm membrane, and stored at −80°C until use. When required, vector particles were concentrated by ultracentrifugation (27,000 rpm, 2 h, 4°C).

#### Generation of Stably Transduced GL261 Cells

To generate stably transduced GL261 cells, cells were infected with LV-RFPL4b/KR or LV-RFPL4b/iGluSnFR and selected using puromycin (2 µg/mL). Stable expression of KR or KR/iGluSnFR was confirmed prior to downstream assays.

### Oncolytic HSV-1 Construction, Reconstitution and Titration

Recombinant oncolytic HSV-1 (oHSV1) variants were generated using bacterial artificial chromosome (BAC) mutagenesis, as previously described [51].

A BAC containing the full genome of HSV-1 strain 17+, with γ34.5 deletions and an FLuc expression cassette inserted into the UL55-UL56 intergenic region, was kindly provided by Beate Sodeik (Hannover Medical School, Germany). This BAC was further modified to:

1. Introduce a Us12 deletion, mirroring the deletion present in T-VEC.
2. Incorporate multiple mir124 target sequences to enhance neuroattenuation.
3. Replace the FLuc cassette with a KillerRed (KR) expression cassette, where KR was targeted to the mitochondria via a COX8 presequence, or alternatively, with mCherry, both under the control of the human CMV (hCMV) promoter.
4. Generate oHSV1-mIL12-aPD1-KR, incorporating a second expression cassette encoding murine IL-12 (mIL12) and an anti-PD1 single-chain antibody (aPD1) into the γ34.5 locus, under the control of the RFPL4b promoter.

mIL12 subunits were linked by a GGGGSGGGGS flexible linker.

aPD1 heavy and light chains were linked by a P2A autoproteolytic peptide.

mIL12 and aPD1 expression cassettes were separated by a T2A autoproteolytic peptide.

All intermediate plasmids used for BAC mutagenesis and the final bacmids were verified via restriction analysis and/or sequencing.

Recombinant oHSV1s were reconstituted by transfecting HEK-293T cells with the corresponding bacmid DNA, followed by viral amplification and titration in highly permissive Vero cells, as previously described [52]. The same titration assay was adopted for the quantification of infectious viral particles released in cell culture supernatants, when requested.

### Monocyte Isolation, Infection, and Functional Assays

#### Human Monocyte Isolation

Primary human monocytes were isolated from buffy coats obtained from healthy donors, provided by the Transfusion Center of the University Hospital of Padua, Italy. Peripheral blood mononuclear cells (PBMCs) were first separated by density gradient centrifugation using Ficoll-Paque® Plus (GE17-1440-03, Sigma-Aldrich), following the manufacturer’s instructions. CD14+ monocytes were then purified from the PBMC fraction using the EasySep™ Human CD14 Positive Selection Kit II (StemCell Technologies), according to the provided protocol.

For select experiments, monocytes were cultured in cancer cell-conditioned medium derived from subconfluent U87-MG and LN229 cells, which were grown in RPMI 1640 medium supplemented with 10% FBS and 1% Penicillin-Streptomycin for 24 hours before harvesting the supernatant.

#### Murine Monocyte Isolation

Primary murine monocytes were isolated from bone marrow and spleen of 12-week-old male C57BL/6J mice using the EasySep™ Mouse Monocyte Isolation Kit (StemCell Technologies). For specific applications, commercially available C57BL/6J mouse bone marrow monocytes (CliniSciences, C57-6271F) were used.

#### Monocyte Infection and Co-Culture Assays

Primary human and murine monocytes were infected with oHSV1 at a multiplicity of infection (MOI) of 3–5 plaque-forming units (PFU)/cell, depending on the experimental conditions.

For co-culture assays, infected monocytes were resuspended in complete culture medium and co-cultured with confluent cancer cells at a 1:1 ratio. Supernatants were collected at fixed time points for viral titration.

#### Migration Assays

Monocyte migration was assessed using 6.5 mm Transwell plates with 5.0 μm pore polycarbonate membrane inserts (Corning, Torino, Italy). Uninfected and oHSV1-infected human monocytes (10⁵ cells) were suspended in 100 µL serum-free OptiMEM (Gibco) and seeded into the upper chamber of the Transwell inserts. The lower chamber contained 600 µL of either serum-free medium alone or cancer cell-conditioned medium. Transwell plates were incubated for 3 hours at 37°C in a 5% CO₂ atmosphere with 98% humidity. Following incubation, inserts were removed, and migrated cells were stained with CellTracker™ Green Dye (Invitrogen, Thermo Fisher Scientific) following the manufacturer’s protocol. Migrated cells were counted across at least three different 10× microscopy fields.

### GBM Cell Infection with oHSV1

GBM cells were seeded in appropriate multi-well plates, depending on the cell type and experimental requirements: 96-well plates: 1×10^4^–1.5×10^4^ cells/well; 24-well plates: 1×10⁵ – 1.8×10⁵ cells/well; 6-well plates: 2.5×10⁵ – 8.7×10⁵ cells/well.

The following day, cells were infected with recombinant oHSV1 variants at the appropriate multiplicity of infection (MOI) in serum-free DMEM. After 1-hour incubation at 37°C, cells were washed three times with PBS and then maintained in DMEM supplemented with 2% FBS.

Infected cells were observed at various time points using fluorescence or confocal microscopy. When required, supernatants or cellular pellets were collected for downstream assays.

#### Spheroid Infection Assays

GBM spheroids were infected with pre-determined PFU of each oHSV1 variant. The desired viral dose was prepared in serum-free DMEM and added to the Ultra-Low Attachment (ULA) 96-well round-bottom plates (Corning® 7007) containing pre-formed spheroids.

After 1-hour incubation at 37°C, allowing for viral adsorption, the medium was replaced with DMEM supplemented with 10% FBS. Fluorescence microscopy was used in the following days to monitor viral infection and spread within the spheroids.

### GBM Spheroid Infection with oHSV1-Loaded Monocytes

To evaluate monocyte-mediated oHSV1 delivery, U87-MG or GL261 spheroids were co-cultured with oHSV1-mCherry-infected human or murine primary monocytes, respectively.

#### Monocyte Labeling and Infection with oHSV1-mCherry

Freshly isolated monocytes were labeled with CellTracker™ Green by incubating them in serum-free RPMI medium containing 1 µL of dye per 1 mL of solution for 20 minutes at 37°C. Labeled monocytes were pelleted by centrifugation (1200 rpm, 5 minutes), washed twice with 1 mL PBS, and resuspended in serum-free RPMI. Monocytes were then infected with oHSV1-mCherry at a MOI of 3-5 PFU/cell by incubating them for 60 minutes at 37°C. Following infection, cells were pelleted, washed twice with PBS, and resuspended in DMEM supplemented with 10% FBS or PBS, depending on the experiments.

#### Co-Culture with GBM Spheroids

oHSV1-loaded monocytes were added to pre-formed tumor spheroids at a final concentration of 5×10³ monocytes per spheroid. Infection and viral spread were monitored over time using fluorescence microscopy.

### Real-Time PCR Analysis

Real-time PCR (qPCR) was performed on cDNA derived from RNA isolated from cells infected with oHSV1-KR-mIL12-aPD1.

#### RNA Extraction and RT-qPCR for Gene Expression Analysis

Total RNA was extracted using the RNeasy Kit (Qiagen). Reverse transcription and qPCR were performed using AgPath-ID™ One-Step RT-PCR Reagents (Life Technologies), according to the manufacturer’s protocol.

A list of the primers and probes adopted in the study is reported in Table 1.

**Table 1.**
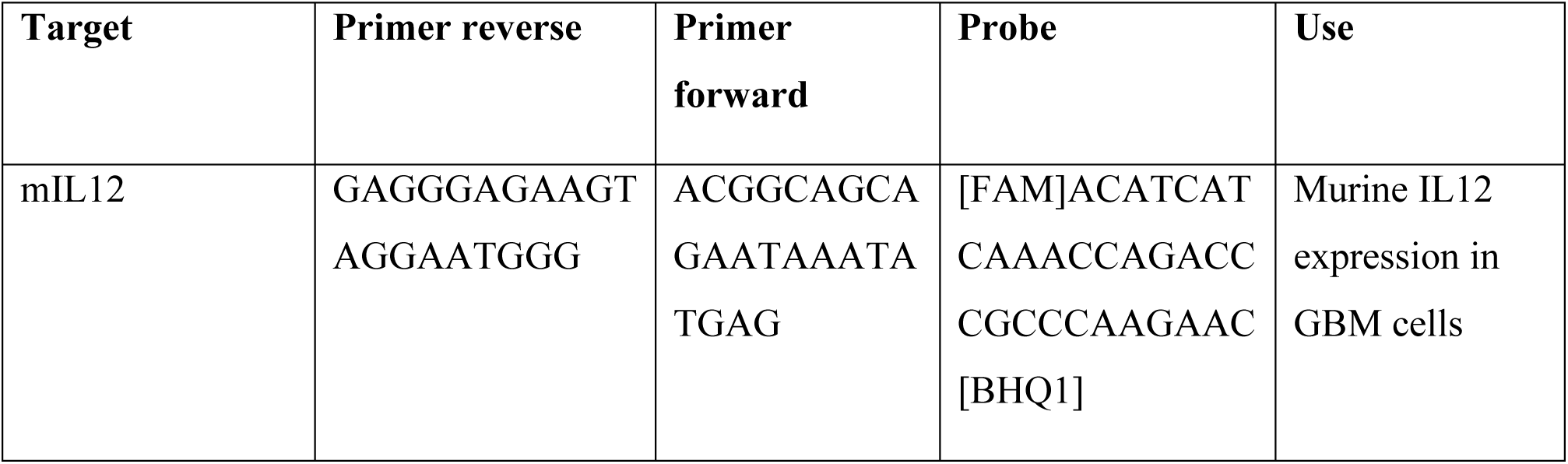
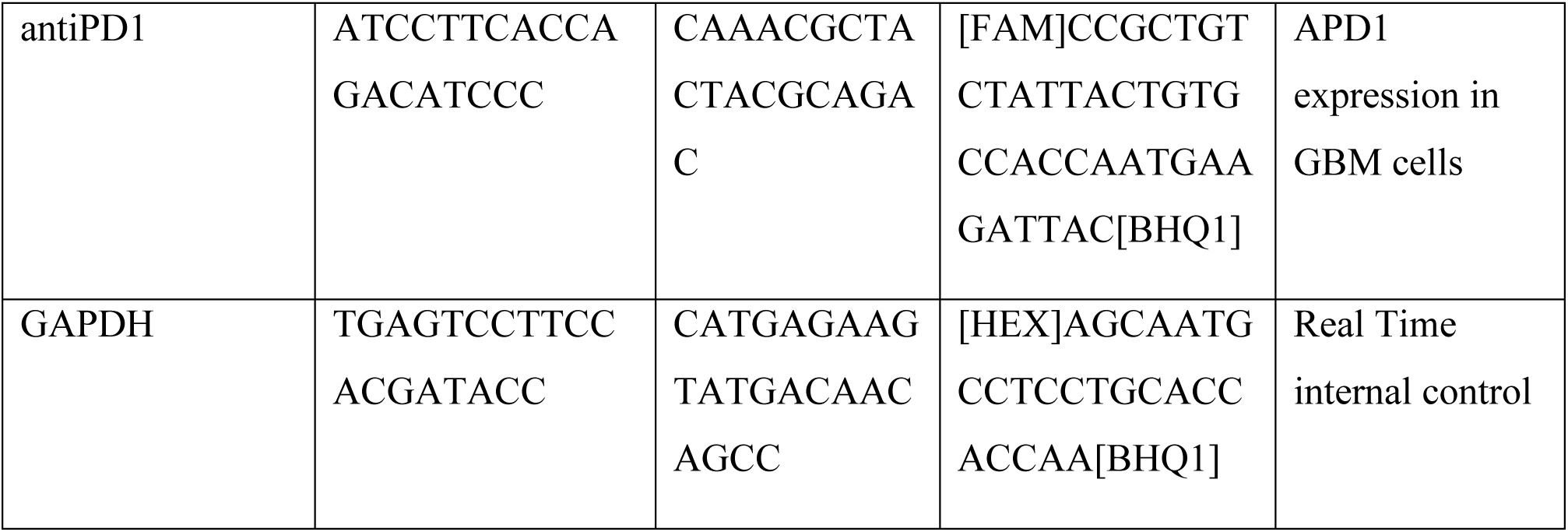
List of primers and probes adopted and their use.

### Evaluation of KR and mIL12 Expression in oHSV1-Infected Cells

In selected experiments, KR expression was assessed by measuring fluorescence intensity using a Varioskan™ LUX Multimode Microplate Reader (Thermo Fisher Scientific). To this end, GL261 cells were seeded in a black 96-well plate (10^4^ cells/well). The following day, cells were infected with oHSV1-KR at an MOI of 1, 10, or 100 PFU/cell. After infection, cells were maintained in FluoroBrite DMEM supplemented with 10% FBS and 4 mM glutamine. KR fluorescence was recorded at different time points post-infection, using an excitation wavelength of 585 nm and an emission wavelength of 610 nm.

To quantify the amount of mIL12 released into the supernatant, 10⁴ Vero cells were seeded in each well of a 96-well plate. The following day, cells were infected with oHSV1-KR-mIL12-aPD1 at an MOI of 0.1 PFU/cell. At 72 hours post-infection, supernatants were collected and analyzed using a Mouse IL-12 p70 ELISA Kit (Biotechne), following the manufacturer’s protocol.

### Photoactivation of KR *in vitro*

GL261-KR cells were seeded in a TC-treated 96-well plate (104 cells/well) in Dulbecco’s Modified Eagle Medium (DMEM) supplemented with 1% penicillin/streptomycin and 10% fetal bovine serum (FBS). The following day, 15µL of polarity-Sensitive Indicator of Viability & Apoptosis (pSIVA)-IANBD (Biorad) were added to each well. pSIVA-IANBD is a green, fluorescent probe which binds extracellular phosphatidylserine, thus serving as an indicator of early apoptosis. Cells were then photoactivated using the 561nm laser of a confocal microscope (Nikon Ti Eclipse) at 100% power under a 20x objective, until complete bleaching of the KillerRed protein fluorescence was observed (75 minutes). pSIVA fluorescence was monitored using the 488 nm laser of the confocal microscope at 5% power. After irradiation was completed, cellular fluorescence was further monitored 1 hour and 3 hours later. Non-irradiated cells were monitored at the same time points without the 75 minutes irradiation step.

GL261 cells were seeded in a 96 well plate (5×10^3^ cells/well). Twenty-four h later, cells were infected with oHSV1-KR at the MOI of 10 PFU/cell or left uninfected as a control. Forty-eight hours later, three wells of infected cells were irradiated for 30 minutes with a 565 nm led. Three wells of infected cells were not irradiated as a control. Twenty-four hours later, cells were detached, and their viability was evaluated by trypan blue exclusion assay, as previously described [53].

### Animal Strains, Housing, Care, and Ethics

C57BL/6J mice were provided and bred by the Consiglio Nazionale delle Ricerche-European Mouse Mutant Archive (CNR-EMMA)-Infrafrontier specific pathogen-free (SPF) unit (Monterotondo Scalo, Rome, Italy) and by Envigo RMS (Bresso, Milan, Italy). After weaning, mice were housed in litters of the same sex, with three to five animals per cage in ventilated cages maintained at a temperature of 20 ± 2°C and a relative humidity of 55 ± 15%. The air within the facility was exchanged 12 to 15 times per hour, and mice were kept under a 12-hour light/dark cycle. Cages were furnished with certified dust-free wood bedding (Scobis One, Mucedola, Settimo Milanese, Milan, Italy), and animals were fed a standardized diet (4RFN and Emma 23, Mucedola) with ad libitum access to chlorinated, filtered water.

Treated and control mice, both male and female, were individually housed and monitored daily for health and well-being. All experimental procedures involving animals were reviewed and approved by the local animal welfare oversight bodies and were conducted under the direct supervision of the CNR-IBBC/Infrafrontier-Animal Welfare and Ethical Review Body (AWERB). The study was performed in accordance with general guidelines for animal experimentation, as approved by the Italian Ministry of Health, in compliance with Legislative Decree 26/2014 (Project License 335/2024-PR), which transposes Directive 2010/63/EU on the protection of animals used for scientific purposes. In addition, all experimental procedures adhered to the NIH Guide for the Care and Use of Laboratory Animals and followed the recommendations outlined in both the ARRIVE and PREPARE guidelines [54, 55].

### Biodistribution Analysis of oHSV1-mCherry in Healthy Animals

To assess the biodistribution of oHSV1-mCherry, intra-striatal injections were performed in adult C57BL/6J mice using stereotaxic coordinates from bregma: AP = +0.86 mm, ML = −1.8 mm, DV = −3 mm. Each mouse received 1 μL of oHSV1-mCherry (1.8×10⁵ PFU), following standard surgical procedures.

Post-injection, animals were weighed and monitored daily for general health status, including body weight changes and neurological signs. Mice were sacrificed at 7- or 14-days post-infection, and tissues were collected for further analysis of viral distribution.

### In Vivo Syngeneic Intracranial Tumor Models

For tumor implantation, GL261 cells were cultured to 70% confluence, harvested using 0.05% Trypsin at 37°C for 3 minutes, and resuspended in DMEM complete culture medium. Prior to implantation, cells were centrifuged at 1000 rpm for 3 minutes, resuspended in Hanks’ Balanced Salt Solution (HBSS, cat. n. 14025-050, Gibco), and counted using an automated cell counter (Countess, cat. n. C10227, Invitrogen, Thermo Fisher Scientific). The final suspension was prepared in HBSS at the required concentration for intracranial injections.

Mice were anesthetized and positioned in a stereotaxic apparatus (cat. n. 68513, RWD, TX, USA) using ear bars for head stabilization. A 0.5 mm craniotomy was performed at the designated coordinates from bregma, depending on the tumor implantation site. For intracortical injections, cells were injected at AP = −1 mm, ML = +1.5 mm, DV = −1.1 mm, whereas for intrastriatal injections, coordinates were AP = +0.86 mm, ML = +1.8 mm, DV = −3 mm.

A 2 μL suspension containing 3×10⁴ GL261 cells/μL was delivered at a speed of 1 nL/s using beveled glass capillaries with a 100 μm diameter aperture, mounted on a 100 µL NanoFil syringe (World Precision Instruments, WPI, FL, USA) prefilled with mineral oil (cat. n. M5904, Sigma-Aldrich, Merck, MA, USA) and connected to a microinjection pump (PhD Ultra 70-3601 Nanomite Syringe Pump, Harvard Apparatus, MA, USA).

Following cell implantation, the capillary was left in place for 15 minutes and withdrawn slowly to prevent fluid spillover. The craniotomy was then sealed with bee wax, and the wound was sutured to complete the procedure.

### Intratumoral Injection of oHSV1-mCherry or Infected Monocytes

Intratumoral injections of oHSV1-mCherry (1.8×10⁵ PFU) or oHSV1-mCherry-infected monocytes (5×10⁴ infected monocytes), infected as detailed above, were performed one week after GL261 cell implantation. The injections were administered using the same stereotactic coordinates as those used for tumor implantation (see above).

Following treatment, mice were weighed and monitored daily for general health status, including body weight changes and signs of distress. Animals were sacrificed two weeks post-treatment, and tissues were collected for further analysis.

### Immunofluorescence Analysis

Dissected brains were post-fixed overnight at 4°C in 4% paraformaldehyde (PFA, cat. n. P6148, Sigma-Aldrich) prepared in 1× PBS (cat. n. 161-0780, Bio-Rad, CA, USA). Following fixation, tissues were embedded in 2% agarose (cat. n. A0575, Sigma-Aldrich) and serially sectioned into 50 μm coronal slices using a Vibratome (VT1000 S, Leica Biosystems, IL, USA).

For immunostaining, sections were permeabilized with 0.5% Triton X-100 (cat. n. 108603, Merck) in 1× PBS for 1 hour at room temperature. After permeabilization, samples were incubated for 1 hour in blocking buffer, containing 5% normal donkey serum (cat. n. S30, Sigma-Aldrich), 3% bovine serum albumin (BSA; cat. n. A4503, Sigma-Aldrich), and 0.05% Tween-20 (cat. n. 1706531, Bio-Rad) in 1× PBS.

Tissue sections were then incubated overnight at 4°C with primary antibodies diluted in blocking buffer (antibodies listed in **Table 2**). After extensive washing, samples were incubated with fluorophore-conjugated secondary antibodies (listed in **Table 3**). Nuclei were counterstained with 4’,6-diamidino-2-phenylindole (DAPI, cat. n. D1306, Molecular Probes), and slides were mounted using ProLong™ Antifade Mountant (cat. n. P36934, Invitrogen).

**Table 2.** List of primary antibodies used for immunohistochemistry.

| Name | Company | Species | Cat. Number | Dilution |
| --- | --- | --- | --- | --- |
| AlexaFluor 488-anti CD3 | BioLegend | Rat; mAB | 201406 | 1:100 |
| CD68 | Santa Cruz | Mouse; mAB | Sc-20060 | 1:100 |
| mCherry | Proteintech | Rabbit; pAB | 26765-1-AP | 1:800 |
| Gfap | Pharmingen | Mouse; mAB | 556327 | 1:100 |
| Iba1 | Wako Chemicals | Rabbit; pAB | 019-19741 | 1:500 |

Abbreviations: mAB, monoclonal antibody; pAB, polyclonal antibody.

**Table 3.** List of secondary antibodies used for immunohistochemistry.

| Name | Company | Cat. Number | Dilution |
| --- | --- | --- | --- |
| Donkey anti-Rabbit,<br>Alexa Fluor 555 | Thermo Fisher Scientific | A-31572 | 1:800 |
| Donkey anti-Mouse,<br>Alexa Fluor 555 | Thermo Fisher Scientific | A-31570 | 1:800 |
| Goat anti-mouse<br>Alexa Fluor 488 | Thermo Fisher Scientific | A-11029 | 1:800 |
| Donkey anti-Rabbit,<br>Alexa Fluor 488 | Thermo Fisher Scientific | A-21206 | 1:800 |

#### Immunofluorescence image acquisition and analysis

Immunofluorescence images were acquired with a TCS SP5 confocal microscope (Leica Microsystems) equipped with a 40× oil immersion objective (HCX PL Apo, UV optimized, N.A. 1.25, oil, Leica Microsystems) using Leica Application Suite Advanced Fluorescence 2.7.3.9723 (LAS AF) software or with FV1200 confocal microscope (Olympus Corporation) equipped with 40× oil immersion objective (Olympus Corporation, UPLFLN 40×O, N.A. 1.30) and visualized with the FV10-ASW software (version 4.2; Olympus Corporation) [56]. Lower magnification images were acquired with a motorized Leica LMD7000 Microdissection System (Leica Microsystems) using the manufacturer’s imaging software (Leica application suite X, 3.6.0.20104).

#### Quantification of CD3+ T cells

Images of sections containing brain tumors and labelled with anti-CD3 antibody were acquired with the TCS SP5 confocal microscope. Quantification of CD3+ T cells was performed using Fiji/ImageJ by thresholding and the number of cells was divided by the area of the field of view (FOV).

#### Quantification of CD68 immunofluorescence intensity

Images from tumor bearing brain sections were acquired with the LMD7000 Microdissection System using the manufacturer’s image acquisition software. To quantify CD68 immunoreactivity, using Fiji/ImageJ analysis software, for each slice circular regions of interest (ROIs) were selected within the tumor mass and in the contralateral hemisphere and the median fluorescence was computed for each ROI. The fluorescence measured in the tumor region was then normalized to each contralateral fluorescence signal.

#### Quantification of Gfap- and Iba1-positive relative areas

Images on sections containing brain tumors, labelled with anti-Gfap or anti-Iba1-antibodies and acquired with the TCS SP5 confocal microscope were analyzed using Fiji/ImageJ. Gfap- and Iba1- positive pixels were identified by thresholding and their number was divided by that of the corresponding tumoral or peritumoral regions.

## Supplementary Information

## Authors’ contributions

Conceptualization, A.C., D.M., F.M.; methodology, A.C., A.R., V.D., D.M., F.M.; formal analysis, A.R., M.P., V.D.; investigation, A.C., A.R., M.V.F., V.D., A.G.D.O.D.R., M.P., A.Ro., C.D.P., M.T., E.R., D.M., F.M.; resources, A.C., M.T. L.P., F.M.; data curation, A.R., V.D., A.G.D.O.D.R. C.D.P.; writing-original draft preparation, F.M.; writing-review and editing, A.C., V.D., D.M., F.M.; visualization, A.R., V.D., C.D.P., D.M.; supervision, A.C., D.M., and F.M.; project administration, F.M.; funding acquisition, F.M. All authors have read and agreed to the published version of the manuscript.

## Data availability statement

Data are available on request from the authors.

## Acknowledgments

Images in Figure 8A-F were acquired by Chiara Cimmino.

## References

1. Weller, M., et al., Glioma. Nat Rev Dis Primers, 2024. 10(1): p. 33.

2. Yool, A.J. and S. Ramesh, Molecular Targets for Combined Therapeutic Strategies to Limit Glioblastoma Cell Migration and Invasion. Frontiers in Pharmacology, 2020. 11.

3. Shimizu, K., et al., Photodynamic augmentation of oncolytic virus therapy for central nervous system malignancies. Cancer Lett, 2023. 572: p. 216363.

4. Sung, H., et al., Global Cancer Statistics 2020: GLOBOCAN Estimates of Incidence and Mortality Worldwide for 36 Cancers in 185 Countries. CA Cancer J Clin, 2021. 71(3): p. 209–249.

5. Reale, A., et al., Perspectives on immunotherapy via oncolytic viruses. Infect Agent Cancer, 2019. 14: p. 5.

6. Mohr, I., et al., A herpes simplex virus type 1 gamma34.5 second-site suppressor mutant that exhibits enhanced growth in cultured glioblastoma cells is severely attenuated in animals. J Virol, 2001. 75(11): p. 5189–96.

7. Lasner, T.M., et al., Toxicity and neuronal infection of a HSV-1 ICP34.5 mutant in nude mice. J Neurovirol, 1998. 4(1): p. 100–5.

8. Streby, K.A., et al., Intratumoral Injection of HSV1716, an Oncolytic Herpes Virus, Is Safe and Shows Evidence of Immune Response and Viral Replication in Young Cancer Patients. Clin Cancer Res, 2017. 23(14): p. 3566–3574.

9. Conry, R.M., et al., Talimogene laherparepvec: First in class oncolytic virotherapy. Hum Vaccin Immunother, 2018. 14(4): p. 839–846.

10. Wang, P., et al., Recombinant adenovirus expressing ICP47 gene suppresses the ability of dendritic cells by restricting specific T cell responses. Cell Immunol, 2013. 282(2): p. 129–35.

11. Liu, B.L., et al., ICP34.5 deleted herpes simplex virus with enhanced oncolytic, immune stimulating, and anti-tumour properties. Gene Ther, 2003. 10(4): p. 292–303.

12. Bernstock, J.D., et al., The Current Landscape of Oncolytic Herpes Simplex Viruses as Novel Therapies for Brain Malignancies. Viruses, 2021. 13(6).

13. Markert, J.M., et al., Phase Ib trial of mutant herpes simplex virus G207 inoculated pre-and post-tumor resection for recurrent GBM. Mol Ther, 2009. 17(1): p. 199–207.

14. Patel, D.M., et al., Design of a Phase I Clinical Trial to Evaluate M032, a Genetically Engineered HSV-1 Expressing IL-12, in Patients with Recurrent/Progressive Glioblastoma Multiforme, Anaplastic Astrocytoma, or Gliosarcoma. Hum Gene Ther Clin Dev, 2016. 27(2): p. 69–78.

15. Kang, K.D., et al., Safety and Efficacy of Intraventricular Immunovirotherapy with Oncolytic HSV-1 for CNS Cancers. Clin Cancer Res, 2022. 28(24): p. 5419–5430.

16. Reale, A., A. Calistri, and J. Altomonte, Giving Oncolytic Viruses a Free Ride: Carrier Cells for Oncolytic Virotherapy. Pharmaceutics, 2021. 13(12).

17. Chen, Z., et al., Cellular and Molecular Identity of Tumor-Associated Macrophages in Glioblastoma. Cancer Res, 2017. 77(9): p. 2266–2278.

18. Waschbisch, A., et al., Pivotal Role for CD16+ Monocytes in Immune Surveillance of the Central Nervous System. J Immunol, 2016. 196(4): p. 1558–67.

19. Bunuales, M., et al., Evaluation of monocytes as carriers for armed oncolytic adenoviruses in murine and Syrian hamster models of cancer. Hum Gene Ther, 2012. 23(12): p. 1258–68.

20. Peng, K.W., et al., Tumor-associated macrophages infiltrate plasmacytomas and can serve as cell carriers for oncolytic measles virotherapy of disseminated myeloma. Am J Hematol, 2009. 84(7): p. 401–7.

21. Venuti, A., et al., HSV-1\EGFP stimulates miR-146a expression in a NF-kappaB-dependent manner in monocytic THP-1 cells. Sci Rep, 2019. 9(1): p. 5157.

22. Miebach, L., J. Berner, and S. Bekeschus, In ovo model in cancer research and tumor immunology. Front Immunol, 2022. 13: p. 1006064.

23. Krutzke, L., et al., Chorioallantoic Membrane Tumor Model for Evaluating Oncolytic Viruses. Hum Gene Ther, 2020. 31(19-20): p. 1100–1113.

24. Domka, W., et al., Photodynamic therapy in brain cancer: mechanisms, clinical and preclinical studies and therapeutic challenges. Frontiers in Chemistry, 2023. 11.

25. Vermandel, M., et al., Standardized intraoperative 5-ALA photodynamic therapy for newly diagnosed glioblastoma patients: a preliminary analysis of the INDYGO clinical trial. J Neurooncol, 2021. 152(3): p. 501–514.

26. Leroy, H.A., et al., Is interstitial photodynamic therapy for brain tumors ready for clinical practice? A systematic review. Photodiagnosis Photodyn Ther, 2021. 36: p. 102492.

27. Bulina, M.E., et al., A genetically encoded photosensitizer. Nature Biotechnology, 2006. 24(1): p. 95–99.

28. Serebrovskaya, E.O., et al., Targeting cancer cells by using an antireceptor antibody-photosensitizer fusion protein. Proc Natl Acad Sci U S A, 2009. 106(23): p. 9221–5.

29. Takehara, K., et al., Targeted Photodynamic Virotherapy Armed with a Genetically Encoded Photosensitizer. Mol Cancer Ther, 2016. 15(1): p. 199–208.

30. Ahmed, A. and S.W.G. Tait, Targeting immunogenic cell death in cancer. Mol Oncol, 2020. 14(12): p. 2994–3006.

31. McArthur, K. and B.T. Kile, Apoptotic mitochondria prime anti-tumour immunity. Cell Death Discov, 2020. 6(1): p. 98.

32. Picca, A., et al., Cell Death and Inflammation: The Role of Mitochondria in Health and Disease. Cells, 2021. 10(3).

33. Galluzzi, L., et al., Targeting immunogenic cell stress and death for cancer therapy. Nat Rev Drug Discov, 2024. 23(6): p. 445–460.

34. van Vloten, J.P., et al., Critical Interactions between Immunogenic Cancer Cell Death, Oncolytic Viruses, and the Immune System Define the Rational Design of Combination Immunotherapies. J Immunol, 2018. 200(2): p. 450–458.

35. Sasso, E., et al., Replicative conditioning of Herpes simplex type 1 virus by Survivin promoter, combined to ERBB2 retargeting, improves tumour cell-restricted oncolysis. Sci Rep, 2020. 10(1): p. 4307.

36. Chhabra, N. and J. Kennedy, A Review of Cancer Immunotherapy Toxicity: Immune Checkpoint Inhibitors. J Med Toxicol, 2021. 17(4): p. 411–424.

37. Tait Wojno, E.D., C.A. Hunter, and J.S. Stumhofer, The Immunobiology of the Interleukin-12 Family: Room for Discovery. Immunity, 2019. 50(4): p. 851–870.

38. Zhang, T., et al., Talimogene Laherparepvec (T-VEC): A Review of the Recent Advances in Cancer Therapy. J Clin Med, 2023. 12(3).

39. Reale, A., et al., Human Monocytes Are Suitable Carriers for the Delivery of Oncolytic Herpes Simplex Virus Type 1 In Vitro and in a Chicken Embryo Chorioallantoic Membrane Model of Cancer. Int J Mol Sci, 2023. 24(11).

40. Reale, A., et al., An oncolytic HSV-1 vector induces a therapeutic adaptive immune response against glioblastoma. J Transl Med, 2024. 22(1): p. 862.

41. Xu, J., et al., miR-124: A Promising Therapeutic Target for Central Nervous System Injuries and Diseases. Cell Mol Neurobiol, 2022. 42(7): p. 2031–2053.

42. Song, B., et al., Adenovirus-mediated shRNA interference against HSV-1 replication in vitro. J Neurovirol, 2016. 22(6): p. 799–807.

43. Demircan, T., et al., Cellular and Molecular Comparison of Glioblastoma Multiform Cell Lines. Cureus, 2021. 13(6): p. e16043.

44. Verlengia, G., et al., Engineered HSV vector achieves safe long-term transgene expression in the central nervous system. Sci Rep, 2017. 7(1): p. 1507.

45. Pistollato, F., et al., Isolation and expansion of regionally defined human glioblastoma cells in vitro. Curr Protoc Stem Cell Biol, 2011. Chapter 3: p. Unit 3 4.

46. Pistollato, F., et al., Intratumoral hypoxic gradient drives stem cells distribution and MGMT expression in glioblastoma. Stem Cells, 2010. 28(5): p. 851–62.

47. Zingales, V., et al., Development of an Easy-To-Use Microfluidic System to Assess Dynamic Exposure to Mycotoxins in 3D Culture Models: Evaluation of Ochratoxin A and Patulin Cytotoxicity. Foods, 2024. 13(24).

48. Trevisan, M., et al., Human neural progenitor cell models to study the antiviral effects and neuroprotective potential of approved and investigational human cytomegalovirus inhibitors. Antiviral Res, 2024. 223: p. 105816.

49. Peres, C., et al., Commercially derived versatile optical architecture for two-photon STED, wavelength mixing and label-free microscopy. Biomedical Optics Express, 2022. 13(3).

50. Naldini, L., et al., In vivo gene delivery and stable transduction of nondividing cells by a lentiviral vector. Science, 1996. 272(5259): p. 263–7.

51. Tischer, B.K., G.A. Smith, and N. Osterrieder, En passant mutagenesis: a two step markerless red recombination system. Methods Mol Biol, 2010. 634: p. 421–30.

52. Vitiello, A., et al., Simultaneous Expression of Different Therapeutic Genes by Infection with Multiple Oncolytic HSV-1 Vectors. Biomedicines, 2024. 12(7).

53. Strober, W., Trypan Blue Exclusion Test of Cell Viability. Curr Protoc Immunol, 2015. 111: p. A3 B 1–A3 B 3.

54. Danos, O., et al., The ARRIVE guidelines, a welcome improvement to standards for reporting animal research. J Gene Med, 2010. 12(7): p. 559–60.

55. Smith, A.J., et al., PREPARE: guidelines for planning animal research and testing. Lab Anim, 2018. 52(2): p. 135–141.

56. Tettey-Matey, A., et al., A fully human IgG1 antibody targeting connexin 32 extracellular domain blocks CMTX1 hemichannel dysfunction in an in vitro model. Cell Commun Signal, 2024. 22(1): p. 589.

57. Collins, T.J., ImageJ for microscopy. Biotechniques, 2007. 43(1 Suppl): p. 25–30.

